# Connexin-46/50 in a dynamic lipid environment resolved by CryoEM at 1.9 Å

**DOI:** 10.1101/2020.04.14.036384

**Authors:** Jonathan A. Flores, Bassam G. Haddad, Kimberly A. Dolan, Janette B. Myers, Craig C. Yoshioka, Jeremy Copperman, Daniel M. Zuckerman, Steve L. Reichow

## Abstract

Gap junctions establish direct pathways for connected cells and tissues to transfer metabolic and electrical messages^1^. The local lipid environment is known to affect the structure, stability and intercellular channel activity of gap junctions^2-5^; however, the molecular basis for these effects remains unknown. To gain insight toward how gap junctions interact with their local membrane environment, we used lipid nanodisc technology to incorporate native connexin-46/50 (Cx46/50) intercellular channels into a dual lipid membrane system, closely mimicking a native cell-to-cell junction. Structural characterization of Cx46/50 lipid-embedded channels by single particle CryoEM revealed a lipid-induced stabilization to the channel, resulting in a 3D reconstruction at 1.9 Å resolution. Together with all-atom molecular dynamics (MD) simulations and 3D heterogeneity analysis of the ensemble CryoEM data, it is shown that Cx46/50 in turn imparts long-range stabilization to the dynamic local lipid environment that is specific to the extracellular lipid leaflet of the two opposed membranes. In addition, nearly 400 water molecules are resolved in the CryoEM map, localized throughout the intercellular permeation pathway and contributing to the channel architecture. These results illustrate how the aqueous-lipid environment is integrated with the architectural stability, structure and function of gap junction communication channels, and demonstrates the ability of CryoEM to effectively characterize dynamical protein-lipid interactions.

## Main

The connexins are a family of transmembrane proteins (21 isoforms in human) that form intercellular channels for cell-to-cell communication^6^. These intercellular channels establish a ∼1.4 nm pore that couples the cytoplasms of neighboring cells, and enable direct passage of electrical and small molecule signals (such as, ions, second messengers, hormones and metabolites)^7^ and therapeutic agents^8^. 10’s – 1000’s of connexin channels may assemble together to form large hexagonally packed arrays, *a*.*k*.*a*. plaques, known as gap junctions. In this way, gap junctions enable the near instantaneous response of electrical synapses in the brain and heart, and contribute to the long-range signaling and metabolic coupling of most tissues. Because of these fundamental roles, aberrant gap junctional coupling is associated with a variety of human diseases, including blindness, deafness, skin disorders, arrhythmia, stroke and cancers^9-11^.

Gap junction intercellular communication is facilitated by a unique macromolecular architecture, where intercellular channels directly couple the plasma membranes of two neighboring cells. The lipid bilayers of opposing cells are separated by a characteristic gap of ∼3.5 nm^12^, a feature for which these structures were first recognized in electron micrographs of cell sections^5,13^. Furthermore, large-scale gap junctional plaque formation is dependent upon a dense mosaic of protein-lipid interactions. *In vitro* reconstitution studies have established that plaque assembly and intercellular channel function are dependent on the lipid environment^2,14,15^. However, the molecular basis for these effects remain largely unknown, due to the lack of high-resolution structural information within a lipid bilayer.

Here, we present a CryoEM structure of native connexin-46/50 (Cx46/50) intercellular channels stabilized in a dual lipid nanodisc system at 1.9 Å resolution – providing an unprecedented level of detail for this class of membrane channels. These structural results are coupled with all-atom molecular dynamics (MD) simulation studies, which together reveal many new features of the connexin channels. Cx46/50 is shown to have a remarkable influence on the local lipid environment, effectively inducing a phase separation (to the gel state) that is specific to the extracellular lipid leaflet of the two opposed membranes. 3D heterogeneity analysis of the CryoEM data identified multiple lipid configurations that co-exist within the dynamic lattice of stabilized lipids, which is further detailed by MD. In addition, ∼400 water molecules are resolved in the CryoEM map, localized at architectural and functionally important sites. Together this work uncovers previously unrecognized roles of the aqueous-lipid environment in stabilizing the structure and assembly of the gap junctions, and suggest Cx46/50 plays an important role in shaping the properties of local membrane environment.

### Structural overview of connexin-46/50 in a dual lipid bilayer

Native (heteromeric/heterotypic) connexin-46/50 intercellular channels were purified from mammalian lens tissue (obtained from sheep), as previously described^16^. Freshly purified channels were reconstituted into self-assembling lipid nanodiscs containing pure dimyristoyl phosphatidylcholine (DMPC) at room temperature (∼25° C), supported by the membrane scaffold protein MSP1E1^17^ (see Methods). Under optimized conditions, the reconstitution resulted in a monodispersed population of intercellular channels embedded into a pair of lipid-nanodiscs, as assessed by size-exclusion chromatography and negative stain EM (Extended Data Fig. 1).

Structure determination by high-resolution single particle CryoEM resulted in a high-quality 3D reconstruction, with an overall resolution of 1.9 Å (gold-standard FSC) (Fig. 1a,b, Extended Data Fig. 2,3 and Supplemental Movie 1). The quality of the CryoEM map allowed for detailed stereo-chemical structural refinement of both Cx46 and Cx50 (Fig. 1b, Extended Data Table 1 and Extended Data Fig. 3). The heteromeric pattern(s) of Cx46/50 co-assembly remain unresolved, following various attempts at computational image classification (see Methods). Nevertheless, atomic models of both Cx50 and Cx46 isoforms were equally well-fit into the D6-symmetrized CryoEM map, reflecting their close sequence and structural similarities, 89% sequence similarity over the structured regions and a resulting 0.16 Å backbone r.m.s.d. (see Methods and Extended Data Fig. 3 for details and limitations regarding the heterogeneity of the natively isolated specimen).

**Figure 1.**
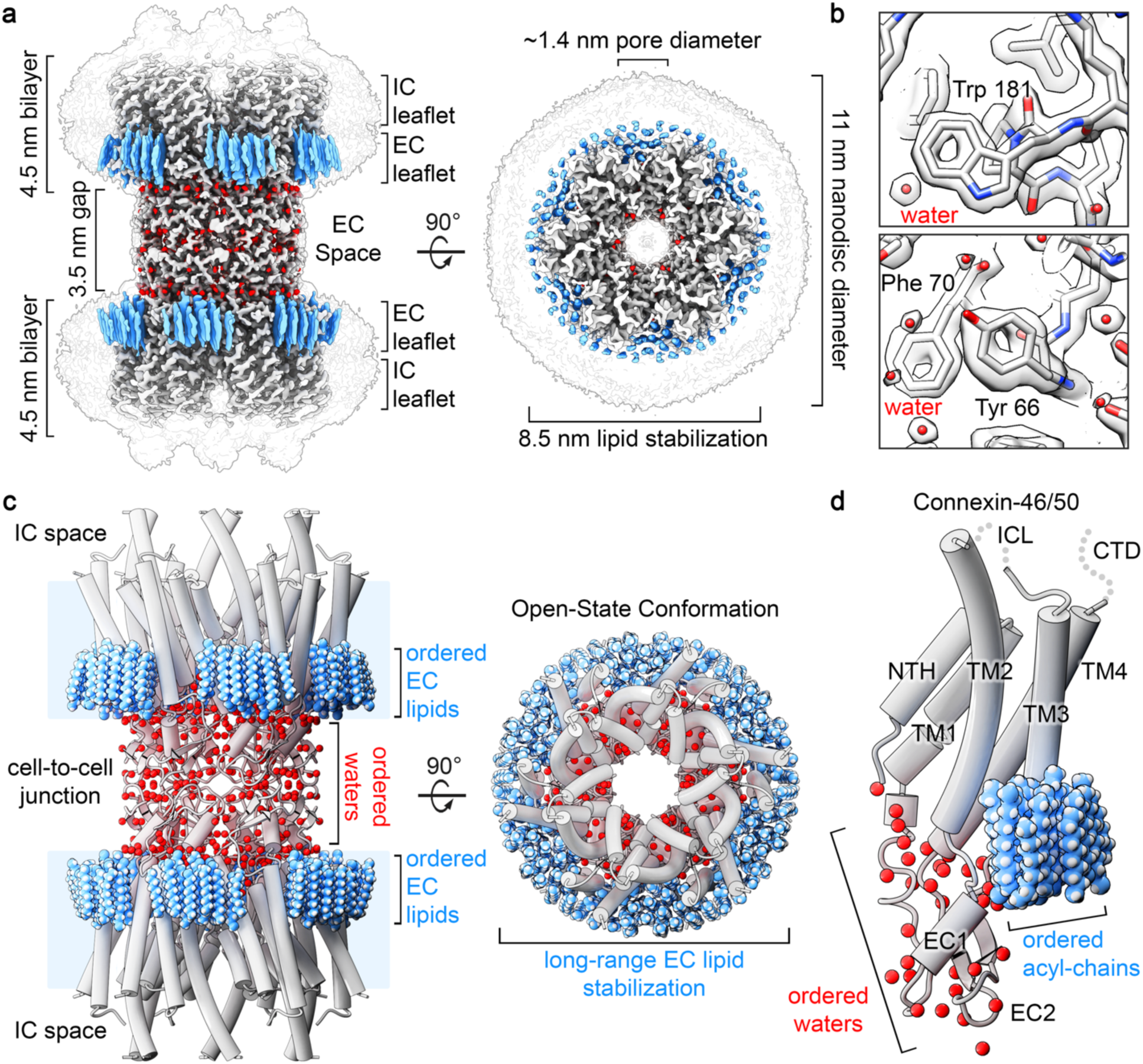
Structure of Connexin-46/50 in lipid nanodiscs by CryoEM. **a)** CryoEM 3D reconstruction of Cx46/50 (white) in an open state conformation, with resolved lipid acyl-chains (blue) and water molecules (red). Transparent silhouette displays the map at low-contour to illustrate the dimensions of the lipid nanodisc densities, with intracellular (IC) and extracellular (EC) lipid leaflets indicated. **b)** Zoom views of the CryoEM map and fitted atomic models, showing high-resolution features observed at 1.9 Å resolution. **c)** Model of Cx46/50 (cylinder representation) with extracellular (EC) lipids and ordered water molecules displayed (spheres). **d)** Cx46/50 monomer, and 15 bound lipids and 33 waters associated with each subunit. Domains labeled for transmembrane helices (TM1-4), extracellular loops (EC1-2) and n-terminal helix (NTH). The intracellular loop (ICL) and c-terminal domain (CTD) are not resolved, indicated by dotted lines.

Cx46/50 is captured in the stabilized open-state, as previously described^16^ (backbone Cα r.m.s.d. = 0.49–0.56 Å and Extended Data Fig. 5), and exposes many new features of the connexin channels that are detailed below. Intercellular channels are constructed by a dodecameric (12-mer) assembly, with six subunits assembled into ‘hemi-channels’ that dock together through extracellular domains, resulting in a continuous ∼1.4 nm pore for intercellular permeation (Fig. 1a,c and Supplemental Movie 2). The distance separating the two lipid nanodisc densities is ∼3.5 nm (Fig. 1a,c), matching that observed by x-ray diffraction on native gap junctional plaques^12^.

Each monomer consists of four transmembrane helices (TM1-4), two extracellular loops (EC1-2) that form the sites of docking interaction and an amphipathic n-terminal helix (NTH), implicated in channel selectivity/gating, is well resolved in the stabilized open-state, as previously described^16^ (Fig. 1c,d and Supplemental Movie 3). However, the significant enhancement in resolution allowed for detailed refinement of sidechain conformations and notable improvement in precision at functional sites, including the NTH domain and the EC1/2 docking sites (Extended Data Fig. 3,5). Furthermore, the quality of the CryoEM map allowed for modeling previously un-resolved regions of TM2 and TM3, which effectively extend the cytoplasmic vestibule of the channel by ∼20 Å, as compared to our previous model (Fig. 1c,d), significantly augmenting the electrostatic environment of the pore entrance (Extended Data Fig. 5). The intracellular loop (ICL) and c-terminal domain (CTD) remain unresolved, presumably due to intrinsic disorder of these regulatory domains^16,18,19^.

Perhaps the most remarkable features of the CryoEM map, however, are the non-protein components of the cell-to-cell junction that are now resolved. A bouquet of 15 ordered lipid acyl-chains is held in place by each of the 12 connexin subunits, which appear to buttress the channel assembly by filling a cavity formed at the lateral subunit interfaces (Fig. 1a,c,d; *blue*). Surprisingly, acyl-chain densities are observed well beyond the first layer of annular lipids that directly interact with the TM domains (primarily TM4 and TM3 of a neighboring subunit) (Fig. 1c,d and Extended Data Fig. 6), suggesting Cx46/50 has a long-range effect on the stability and biophysical properties of the membrane. Remarkably, all of the resolved lipid densities in the CryoEM map are specifically localized to the extracellular leaflet of the bilayer, indicating a selective interaction with the local lipid environment.

In addition to stabilized lipids, 396 ordered water molecules are resolved throughout the channel (33 waters per subunit) (Fig. 1,2; *red* and Extended Data Fig. 6). Waters are found at both solvent accessible and buried sites within the core of the channel, apparently contributing to the permeation pathway and structural integrity of the channel assembly (Fig. 1,2). The assignment of water densities was supported by all-atom equilibrium MD simulations conducted in the presence of explicit water and 150mM NaCl or KCl (see Methods and Extended Data Fig. 7,8). There was no clear evidence that the resolved solvent sites correlated with low-affinity ion binding sites observed by MD (not shown). In the following sections, we describe these newly resolved features in further detail and discuss their potential structural and functional roles.

**Figure 2.**
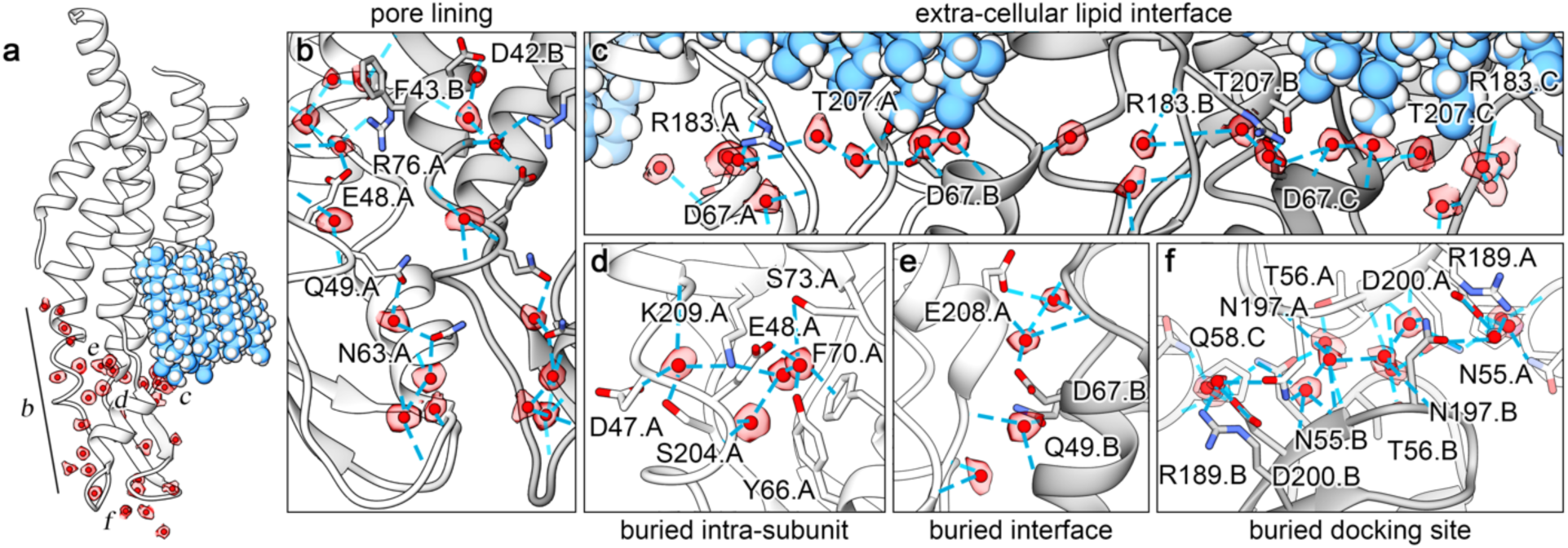
Ordered water molecules resolved in Cx46/50 by CryoEM. **a)** Cx46/50 subunit with segmented CryoEM density of waters overlaid in transparency (colored as in Fig. 1). Labels in panel a indicate position of the various zoom views, presented in panels **b–f**, showing water molecules bound to **b)** pore-lining sites, **c)** extra-cellular lipid interface, **d)** buried intra-subunit sites, **e)** buried subunit interface sites, and **f)** buried cell-to-cell docking sites. In panels b-f, amino acids sidechains forming hydrogen bonds to water are displayed (blue dotted lines) and labeled using Cx50 numbering.

### Stabilized waters contribute to the permeation pathway and core architecture of Cx46/50

Gap junctions establish aqueous pathways that allow a variety of cytosolic substrates, less than ∼1 kDa in size, to permeate from cell to cell^20^. The permeation pathway is established by the pore-lining NTH domain, TM1/2 and EC1 domains (Fig. 1c,d). Within the channel pore of Cx46/50, there are 108 waters bound at solvent-exposed sites (9 per subunit). Pore-bound waters localize to regions of the EC1 domain and TM1 parahelix, and mediate an extensive network of h-bonding interactions, involving, D42, F43 (π bonding), E48, Q49, N63 and R76 in Cx50 (positions 42 and 43 are Glu in Cx46), and several protein backbone interactions (Fig. 2a,b and Extended Data Fig. 6).

EC1 and the TM1 parahelix contribute to the selectivity, conductance and slow (loop) voltage-gating mechanisms of Cx46/50^16,21-23^ and other connexins^24-29^, and are implicated in Ca^2+^-regulation in Cx26 by X-ray crystallography^19^, MD studies^27,30^, and by functional mutation studies of Cx46^30^. As such, these pore-lining waters may functionally contribute to these mechanisms, for example, by orienting or extending the hydrogen-bonding potential of amino-acid sidechains involved in the coordination of substrates (or regulatory ions), buffering the electrostatic properties of the channel pore, or integrating the electrostatic network that is proposed to couple EC1/TM1 to the fast (NTH) voltage-gating domain^31,32^.

On the extracellular surface of the channel, symmetry-related rings of tightly bound water molecules are organized at the extracellular aqueous-lipid boundary (Fig.1a,c and Fig. 2a,c). In the ensemble CryoEM map, the PC lipid head groups are not resolved (due to local disorder described in the following sections). Nevertheless, these stabilized rings of water are nominally positioned at the acyl-headgroup boundary of the extracellular lipid leaflet. These waters are stabilized by hydrogen bonds with EC1/2 residues (D67, R183/Q171 and T207/T195 in Cx50/Cx46, respectively) and expected to be further coordinated through non-specific interactions with the phospho-glycerol backbone of the extracellular PC lipids (Fig. 2c; and discussed below).

The EC1/2 domains appear to be the most well-ordered region of the channel, as reflected by local-resolution of the CryoEM density map (Extended Data Fig. 3) and root-mean-square-fluctuation (r.m.s.f.) analysis of MD-trajectories (Extended Data Fig. 7) This high-degree of stability reflects the important functional role of the EC1/2 domains in maintaining an electro-chemical seal at the cell-to-cell junction. Several clusters of water molecules are found buried a sites located both within and between the EC1/2 domains of individual subunits (Fig. 2a, d-f). A cluster of four waters are buried within the EC1/2 domains is coordinated by residues D47, E48, Y66, F70 (π bonding), S73, S204/S192 and K209/K197, in Cx50/46 respectively (Fig. 2d). Four additional waters are buried at the lateral EC domain interface formed by neighboring subunits, primarily coordinated by hydrogen bonding interactions with the peptide backbone and sidechains of Q49, D67 and E208/E196, in Cx50/Cx46 respectively (Fig. 2e). The degree of coordination of these buried waters suggest they contribute to the architectural integrity of EC1/2 docking domains, and may in part explain why deleterious mutations at D47, E48 and D67 in Cx50 linked to cataract formation disrupt junctional coupling and/or biogenesis^33-35^.

The EC1/EC2 domains also play important roles in establishing the specificity of hemi-channel docking interactions formed between different connexin isoforms, and the ability to establish so-called homotypic or heterotypic channels^36-38^. Elucidating the determinants of hemi-channel recognition is therefore critical to understanding the principles dictating cell-type specificity of gap junctional coupling^39^. It has been proposed that isoform-specific hydrogen bonding patterns that bridge the EC1/EC2 interface govern hemi-channel docking compatibility^38,40^. Contributing to this bridging site in Cx46/50 is a cluster of 12 water molecules (per subunit pair) that are deeply integrated within a dense network of hydrogen bonds between EC1/EC2 residues of opposed subunits (Fig. 2f). At the center of this network is the highly conserved K/R-N-D motif found in EC2 of Group I compatible connexins (including Cx50, Cx46, Cx32 and Cx26). Genetic mutations of this motif in Cx46/50 are linked to congenital cataracts^16,41^, as well as other genetic disorders (*e*.*g*., Charcot-Marie-Tooth disease^42^ and non-syndromic deafness^43^), when mutated in other Group I connexins. These observations suggest interfacial waters may play previously unappreciated and functionally important roles in establishing the structural integrity of the intercellular channel and contribute to the specificity of hemi-channel docking interactions involved in regulating the formation of intercellular communication pathways.

### Cx46/50 induces long-range ordering at the extracellular lipid leaflet

The degree of long-range stabilization to the local lipid environment observed in the Cx46/50 nanodisc reconstruction, extending several solvent layers away from the protein, is (to our knowledge) unprecedented. DMPC was selected as a model lipid because of the high PC content of mammalian (sheep) lens^44^, and reconstitution studies show DMPC produces Cx46/50 assemblies that are indistinguishable from those formed with native lipids^14,45^. Due to its complete saturation DMPC has a relatively high phase-transition (*i*.*e*., melting) temperature (T_m_) compared to other biological lipids (T_m_ ∼24° C in pure lipid vesicles^46^). This value is close to the temperature at which reconstitution was performed (∼25° C, room temperature). However, in nanodiscs the melting temperature of DMPC is reportedly higher (∼28° C), due to compartmentalization effects by the MSP scaffold^47^. Nevertheless, the specific localization of stabilized lipids to the extracellular leaflets observed by CryoEM (and also by MD studies, described below) suggested long-range lipid stabilization is induced through interactions with Cx46/50 (Fig 1, 3a).

**Figure 3.**
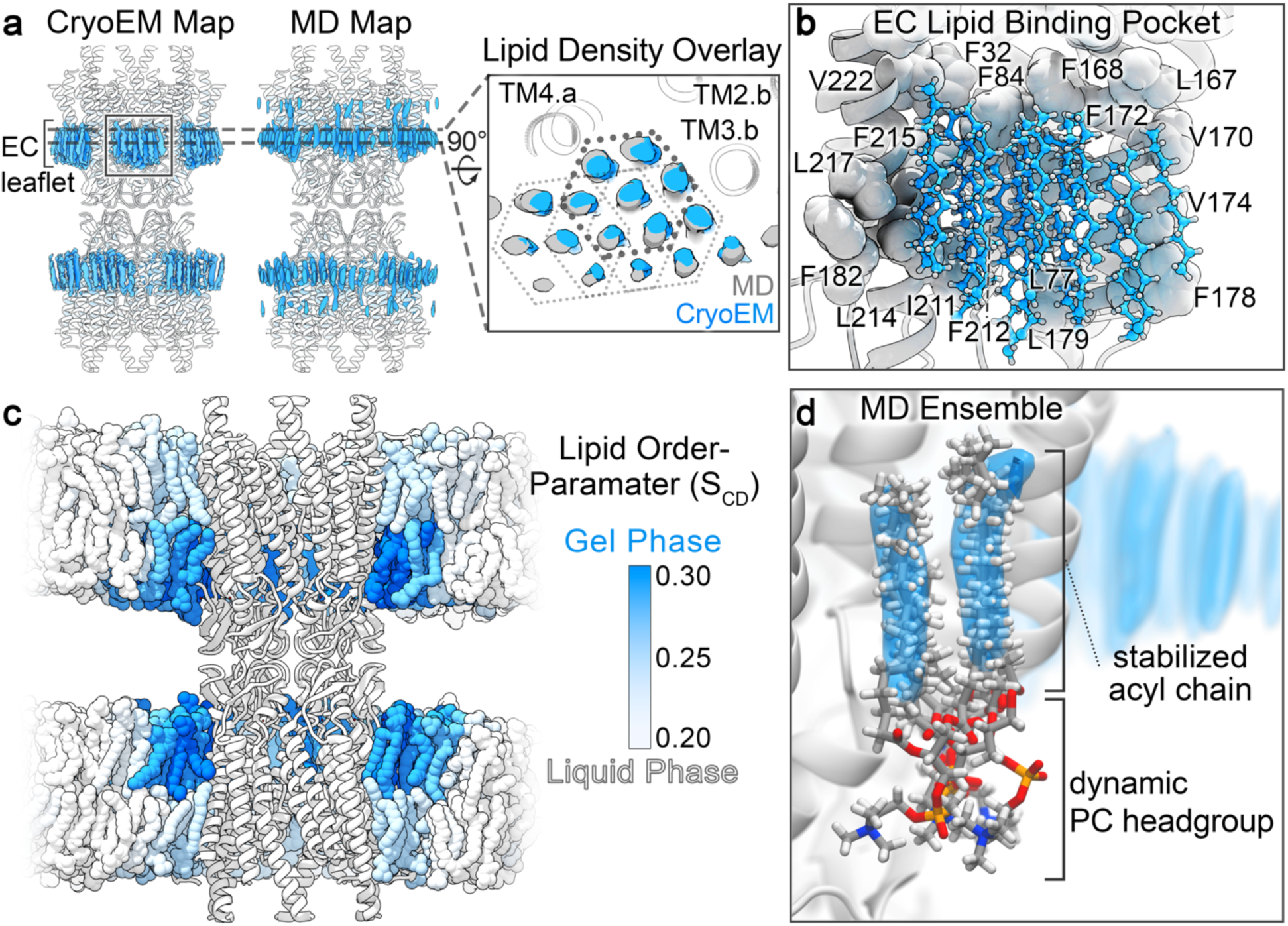
Cx46/50 induces a local phase-separation to the extra-cellular lipid leaflet. **a)** Comparison of acyl-lipid density maps (blue) obtained by CryoEM and time-averaged all-atom MD-simulation, overlaid onto the Cx46/50 ribbon structure (white). *Inset*, shows a slice-view (rotated 90°) of overlaid acyl-lipid densities by CryoEM (blue) and MD-simulation (grey). The hexagonal packing pattern of acyl-chains is indicated (solid and dotted lines), and TM helices interacting with lipid are labeled. Different subunits are indicated by suffix (*a* or *b*). **b)** Zoom-view of the acyl-lipid binding pocket, with lipid-binding residues displayed (spheres) and labeled (Cx50 numbering). **c)** MD-snapshot of Cx50 in phosphatidylcholine (PC) lipid bilayers, with time-averaged lipid order parameter (S_CD_) for each lipid indicated by shading (blue = 0.30 to white = 0.20). **d)** Zoom view, showing an ensemble super-positioning of symmetry-related lipids obtained by MD-simulation (displayed as all atom representation) occupying the MD-based lipid acyl-chain density map (blue).

To gain further insight into the lipid-stabilization observed by CryoEM, we analyzed time-averaged densities of DMPC acyl-chain positions obtained by unbiased all-atom MD simulations for both Cx50 and Cx46, conducted at 37° C, where the starting positions of DMPC molecules had been randomly placed into a 15.4 x 15.4 nm lipid bilayer (see Methods and Extended Data Fig. 7a). Following equilibration, the resulting acyl-lipid density profiles displayed remarkable similarity to what was resolved by CryoEM (Fig. 3a). In both cases, lipids within the extracellular leaflets are specifically stabilized, as compared to the intracellular lipid leaflet (Fig. 3a). Furthermore, the resolved clusters of acyl-chain densities obtained by MD display the same hexagonal packing pattern that extends 3-4 orders beyond the annular shell, as observed by CryoEM (Fig. 3a, *inset*).

The corroborating results obtained by MD imply that the lipid stabilization observed by CryoEM is specifically induced by structural features of the Cx46/50 TM domains, and not an artifact of the nanodisc. Each cluster of lipids is bound by a shallow pocket of hydrophobic and aromatic residues, displayed by TM2/3 and TM4 of adjacent subunits (Fig. 3a,b). A cleft, rich in aromatic sidechains (formed by F32, F84, L167/L155 and F168/F156 in Cx50/C46, respectively) intercalates into the bilayer, appearing to bisect the extracellular leaflet from the more disordered intracellular leaflet (Fig. 3b). In this way, it appears that the acyl-lipid binding pocket selectively grasps a large bouquet of lipids from the extracellular leaflet, inducing long-range stabilization to the membrane through extensive Van der Waals interaction.

The extended acyl-lipid chain conformation and hexagonal packing adopted by the bouquet of bound lipids are indicative of a quasi phase-transition to the liquid-ordered (or gel-like) state. To obtain a more quantitative assessment of the degree of lipid stabilization, we extracted SN1 and SN2 lipid order parameters (S_CD_) from the MD-simulations, which have been parameterized to fit well to experimental NMR-based order parameters^48^. These results are consistent with the notion that Cx46/50 induces a phase transition from a fluid to a gel-like state that is specific to the extracellular lipid leaflet, as indicated by a shift in order parameters to above ∼0.25^49^ (acyl-chain carbons 4–11; Fig. 3c and Extended Data Fig. 9), which extend ∼10–20 Å from the protein surface, as observed by CryoEM.

Although this degree of stabilization to the local lipid environment is likely to depend on lipid type, the general effects may be functionally important. For example, by contributing to the architectural integrity at the gap junctional interface, partitioning specific types of lipids, or even templating long-range hexagonal packing interactions found in plaque assemblies^45,50^. In this context, it is noteworthy that connexins localize to lipid raft domains^51,52^, which are rich in high T_m_ lipids (e.g., sphingomyelin) and characterized as forming a liquid-ordered state.

### Annular PC lipids adopt a dynamic ensemble of conformational and configurational states

Another notable feature of the lipid densities observed in the CryoEM map is that PC head groups are not observed, despite sufficient resolution to expect such features (Fig. 1, 3a). Super-positioning of representative lipid conformations obtained by MD show that, although the annular lipid acyl-chains were relatively well ordered and superimpose, their corresponding head groups remain conformationally dynamic and/or heterogeneously positioned (Fig. 3d and Supplemental Movie 4). Such behavior would rationalize the lack of resolvability in the averaged CryoEM density map. In an attempt to resolve this heterogeneity, we conducted 3D-classification analysis on the ensemble CryoEM data (Methods and Extended Data Fig. 4), which resulted in three distinct 3D reconstructions resolved at resolutions of ∼2.5 Å (gold-standard FSC) (Fig. 4a-c and Extended Data Fig. 4).

**Figure 4.**
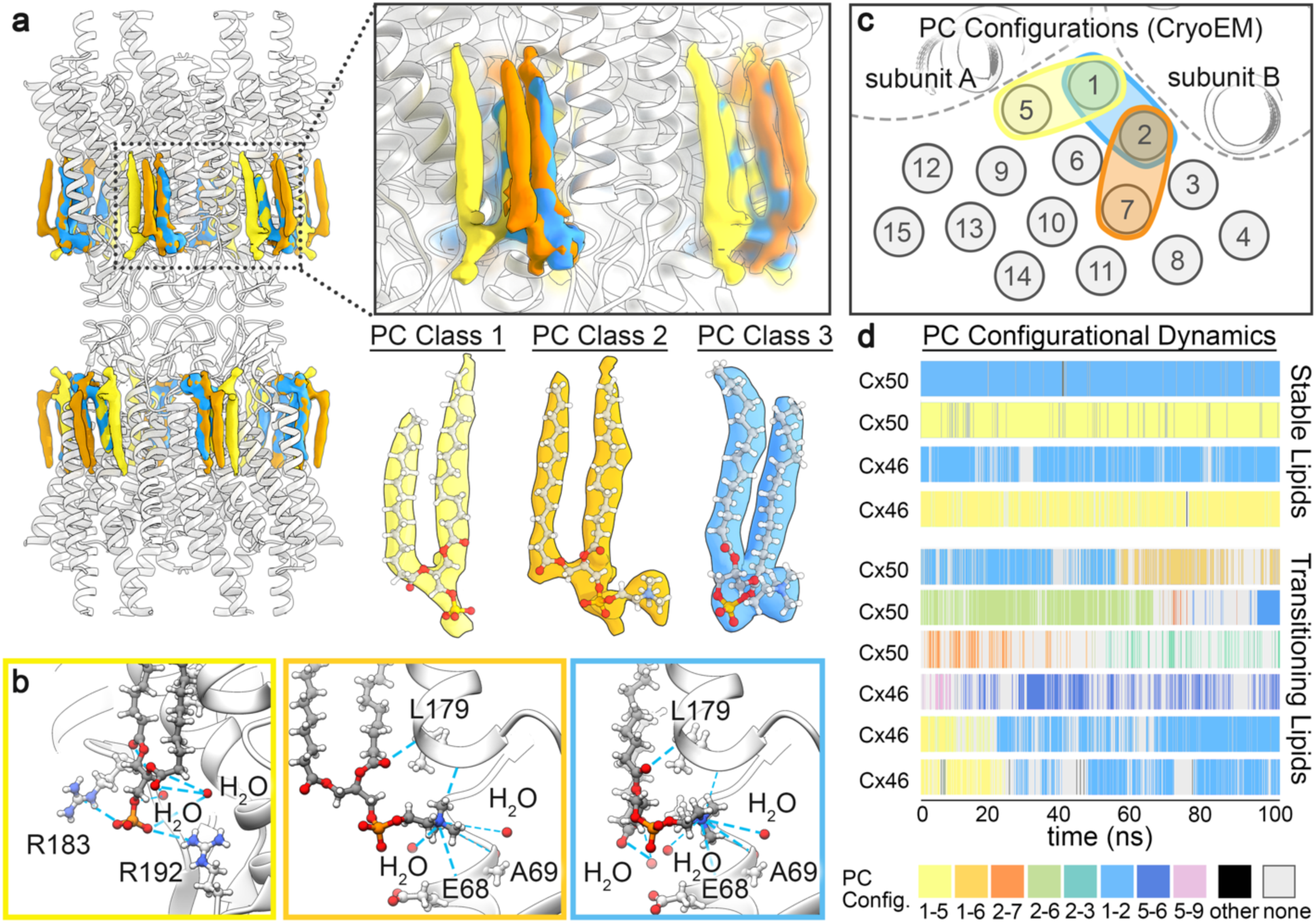
PC lipid configurational heterogeneity and dynamics resolved by CryoEM and MD. **a)** Segmented phosphatidylcholine (PC) density maps obtained by CryoEM 3D heterogeneity analysis and classification (PC Class 1 – yellow, PC Class 2 – orange, PC Class 3 – blue). *Insets*, show a zoom-view displaying the overlapping features of resolved lipid configurations, and segmented densities with fitted atomic-models obtained from the three PC classes. CryoEM density for all other non-unique acyl-lipid chains, with unresolved head groups, have been omitted for clarity. **b)** Zoom-view, showing Cx50 hydrogen bond interactions (blue dotted line) between the PC lipid headgroup and phospho-glycerol backbone. Interacting amino-acids and stabilized water molecules are labeled. Yellow box – PC Class 1, orange box – PC Class 2, blue box – PC Class 3. **c)** Illustration, showing acyl-chain positions and configurational assignments resolved by CryoEM (represented as grey circles and numbered 1 – 15). **d)** PC configurational classification and dwell times obtained by all-atom MD-simulation, showing representative populations of stable (non-transitioning) and dynamic (transitioning) lipids. PC configurations were classified by acyl-chain occupancy in densities numbered as in panel c, and colored uniquely (as indicated, bottom of panel d).

In each of the 3D classes, PC head groups and/or phospho-glycerol backbone of individual annular lipids were uniquely resolved (PC Class 1–3, Fig. 4a). The conformational state of Cx46/50 is very similar in all three classes, and essentially indistinguishable from models derived from the ensemble density map (Cα r.m.s.d.’s = 0.24 – 0.34 Å). Notably, structural features of each of these fully-resolved lipids are shared amongst these classes. For example, the SN2 acyl-chain of PC Class 1 overlays with the SN1 chain of PC Class 3 (Fig. 4a,c; *yellow and blue*). Likewise, the SN1 chain of PC Class 2 also overlays with the SN1 chain of PC Class 3 (Fig. 4a,c; *orange and blue*). This suggested that multiple, overlapping, configurational states are capable of supporting the same lattice of acyl-chain positions observed in the CryoEM reconstructions, consistent with non-specific and/or transient binding interactions.

The choline head group of PC Class 1 remained unresolved, however the negatively charged phospho-glycerol backbone is clearly visualized and appears to be stabilized by flanking positively charged arginine residues R183 and R192 in Cx50 (Q171 and R180 in Cx46) (Fig. 4b; *left*), and hydrogen bonding with two water molecules (part of the belt of extracellular waters described above, see Fig. 2c). PC Class 2 and 3 resolve distinct acyl-chain configurations, yet, both of these states share a similar placement of their positively charged choline head groups. Head group placement of these lipids is supported by non-specific hydrogen-bond interactions with backbone carbonyls presented by EC1 and TM3 (involving residues E68, A69 and L179 in Cx50; position 68 is an Arg in Cx46) and a buried water molecule (Fig. 4b; *center and right*). The phospho-glycerol backbones of PC Class 2/3 lipids are coordinated by hydrogen-bonding to local waters and the backbone amide of L179 in Cx50 (L167 in Cx46). Remarkably, the SN2 acyl chain and glycerol backbone of the PC Class 2 lipid is completely resolved, despite lacking any direct contact with the Cx46/50 protein interface (Fig. 4a,c).

Collectively, these observations support the notion that while the Cx46/50 acyl-chain interactions appear to be high-affinity, the lipid head group interactions are nonspecific and adopt a variety of configurational/conformational states. This is reinforced by our MD-simulation studies for both Cx50 and Cx46, where mapping of PC arrangements at resolved acyl-chain densities show a variety of configurational states that co-exist within the dodecameric assembly. Furthermore, during the timescale of the simulations (100 ns), time-resolved PC configurations could be classified as being either stable or dynamically transitioning between multiple configurational states (Fig. 4d and Supplemental Movie 5,6). Notably, the most stable (yet overlapping) configurations (*e*.*g*., 1-2 and 1-5 configurations) are the same as those resolved by CryoEM 3D classification (Fig. 4a,c,d; *blue and yellow respectively*). Yet, other lipid trajectories were observed interconverting between these same configurations over this relatively short time-scale (Fig. 4d). The degree of configurational preference diminishes beyond the first two solvent shells, presumably due to the loss of energetic influence induced by protein interactions (Fig. 4d), and reflect the randomized head group arrangements expected of a bulk lipid population. Taken together, these data show Cx46/50 stabilizes the dynamic local lipid environment through non-specific interactions with the extracellular leaflet, with multiple configurational PC lipid states existing at the annular interface and effectively captured by CryoEM.

### Concluding Remarks

The structure and function of membrane proteins are deeply integrated with their lipid environment. Our mechanistic understanding of protein-lipid interactions have been largely shaped by high-resolution structures of membrane proteins where specifically bound lipids have been captured at well-defined binding sites^53^. Yet, most interactions made between membrane proteins and their local membrane environment are relatively non-specific and highly dynamic. The mechanistic principles and biophysical consequences underlying such interactions remains poorly understood, as these interactions are typically lost during protein purification, or remain too dynamic to resolve by traditional structural methods. By exploiting the potential of lipid nanodisc technologies coupled with single particle CryoEM and MD simulation, we show that Cx46/50 intercellular communication channels form dynamic interactions with annular lipids. These non-specific interactions have long-range stabilizing effects capable of inducing a phase separation to high T_m_ lipids, which may extend ∼20 Å from the protein surface. These interactions appear selective toward the extracellular leaflet of pure PC membranes, which may have significant consequences on the biomechanical properties and lipid composition of gap junctional domains. In fact, the lack of resolved lipids in the intracellular leaflet may reflect the selectivity at this leaflet toward non-PC lipid types, as suggested for Cx26/32^15^. The methods developed here provide a valuable high-resolution platform for developing our deeper understanding of the specificity and physiological role lipids play in gap junction biology, and how aberrant lipid environments may contribute to connexin-related pathologies. Indeed, the capability of resolving connexin channels beyond the critical threshold of ∼2.0–2.5 Å resolution, the precision desired for structure-based drug design – *e*.*g*., providing detailed stereo-chemical models and placement of architectural water molecules – now opens the door to rational development of selective high-affinity pharmacological tools that are desperately needed in this field to better understand and potentially treat a wide range of connexin-opathies^54^.

## Acknowledgements

We thank Dror Chorev and Carol V. Robinson for help analyzing specimens prepared for CryoEM. We are grateful to the staff at the OHSU Multiscale Microscopy Core and Advanced Computing center, and to the Pacific Northwest Center for CryoEM (supported by NIH Grant U24GM129547) and accessed through EMSL (grid.436923.9). J.A.F. is supported by the National Institutes of Health NRSA (F31-EY030409). K.A.D. is supported by National Institutes of Health BUILD EXITO Program (TL4-GM118965) and Berkeley Molecular Biophysics Training Grant (T32-GM008295). C.C.Y. is supported by OHSU. J.C. and D.M.Z. are supported by the OHSU Center for Spatial Systems Biomedicine, by the National Science Foundation (MCB 1715823) and by the National Institutes of Health (R01-GM115805). S.L.R. is supported by the National Institutes of Health (R35-GM124779).

## Author Contributions

J.A.F, B.G.H and K.A.D. contributed equally. K.A.D. and J.A.F. conducted the protein purification and reconstitution of CryoEM specimens. J.A.F. collected the CryoEM datasets, performed image analysis and atomic modeling. K.A.D., J.B.M., C.C.Y. contributed to image analysis. B.G.H. conducted and analyzed the MD simulations. B.G.H., J.C. and D.M.Z. contributed to the experimental design and analysis of MD simulations. All authors contributed to manuscript preparation. S.L.R. provided overall guidance to the design and execution of the work.

## Author Information

These authors contributed equally: Jonathan A. Flores, Bassam G. Haddad and Kimberly A. Dolan.

## Corresponding Author

Correspondence and requests for materials should be addressed to

## Methods

### MSP expression and purification

A plasmid containing the coding sequence for membrane scaffold protein 1E1 (MSP1E1) was obtained from Addgene^17^ and the protein was expressed and purified as described^55^, with minor modification. Freshly transformed *E. coli* cells (BL21Gold-DE3) were grown in LB medium containing 50 μg mL^-1^ kanamycin at 37° C with shaking (250 rpm). Induction with 0.5 mM Isopropyl β-d-1-thiogalactopyranoside (IPTG) was performed when OD_600_ reached 0.5–0.6, and allowed to express for 3–5 hours post-induction at 37° C. Cells were harvested by centrifugation at 4,000 x *g* for 20 minutes at 4° C, and cell pellets were resuspended in MSP Lysis Buffer (40 mM Tris [pH 7.4], 1% Triton X-100, 1 mM PMSF) at a density of ∼20 mL of Lysis Buffer per Liter of culture. Cell suspensions were flash frozen in liquid nitrogen and stored at –86° C for up to several months.

Frozen cell suspensions were thawed from –86° C storage, supplemented with 1 mM phenylmethylsulfonyl fluoride (PMSF) and lysed by sonication on ice. Crude lysate was cleared by ultra-centrifugation at 146,550 x *g* for 30 minutes at 4° C. The supernatant was filtered (Millipore; 0.22 μm) and applied to a gravity column with 5 mL of HisPur Ni-NTA resin (Thermo Fisher Scientific) prepared in equilibration buffer (40mM Tris [pH 7.4]). MSP-bound resin was washed with 5 column volumes (CV) of equilibration buffer, followed by 5 CVs of each of the following: Triton buffer (40 mM Tris [pH 8.0], 300 mM NaCl, 1% TX-100), Cholate buffer (40 mM Tris [pH 8.0], 300 mM NaCl, 50mM cholate), Imidazole Wash Buffer (40 mM Tris [pH 8.0], 300 mM NaCl, 50 mM imidazole). MSP1E1 was eluted with 3 CVs of Elution Buffer (40 mM Tris [pH 8.0], 300 mM NaCl and 750 mM imidazole). The eluate was filtered (Millipore; 0.22 μm) and applied to a size exclusion chromatography (SEC) column (ENC70; BioRad) equilibrated in 20 mM HEPES (pH 7.4), 150 mM NaCl and 1 mM EDTA using an FPLC (NGC system; BioRad). Peak fractions were monitored by UV_280_, pooled and concentrated to 400-600 μM using a centrifugal device. Final protein concentration was determined by UV absorbance at 280 nm. Samples were aliquoted, flash frozen in liquid nitrogen and stored at –86° C for up to several months.

### Cx46/50 purification and nanodisc reconstitution

Native Cx46/50 intercellular channels were isolated as previously described^16^. Briefly, lamb eyes were obtained from the Wolverine Packers slaughterhouse (Detroit, MI), and the lenses were removed using a surgical blade and stored at –86° C. Gap junction intercellular channels were isolated from the core lens fiber tissue, containing c-terminal truncation variants of Cx46 and Cx50 (*a*.*k*.*a*. MP38)^56-59^. Details of the purification procedure are provided below.

Lenses were thawed from –86° C, core lens fiber cell tissue was dissected from the outer cortical tissue using a surgical blade and stripped core membranes were prepared as described^60-62^. Total protein concentration was determined by BCA (Pierce) and membranes were stored at –86° C, in storage buffer (10 mM Tris [pH 8.0], 2 mM EDTA, 2 mM EGTA) at a total protein concentration of ∼2 mg mL^-1^. Stripped membranes were thawed from –86° C and solubilized with 10 mM Tris (pH 8.0), 2 mM EDTA, 2 mM EGTA, 1% (wt vol^-1^) n-decyl-β-D-maltoside (DM) for 30 minutes at 37° C. Insoluble material was cleared by ultra-centrifugation at 146,550 x *g* for 30 minutes at 4° C. The supernatant was filtered (Millipore; 0.22 μm) and separated by anion-exchange chromatography (UnoQ, BioRad) with buffer A (10 mM Tris [pH 8.0], 2 mM EDTA, 2 mM EGTA, 0.3% DM [wt vol^-1^]). Protein was eluted with a 20 CV gradient of buffer B that additionally contained 500 mM NaCl. Elution peaks containing Cx46/50, as determined by SDS-PAGE, were pooled and applied to a size exclusion chromatography (SEC) column (Superose 6 Increase 10/300 GL; GE Healthcare) equilibrated with SEC buffer (20 mM HEPES [pH 7.4], 150 mM NaCl, 2 mM EDTA, 2 mM EGTA and 0.3% DM [wt vol^-1^]). Peak fractions containing purified Cx46/50 were pooled and concentrated to 5–6 mg mL^-1^ with a centrifugal device (Vivaspin 6; 50-kDa cut-off filter; Sartorius). Protein concentration was determined by UV absorbance at 280 nm. All chromatography steps were performed by FPLC at 4° C.

Freshly purified Cx46/50 was reconstituted into MSP1E1 nanodiscs using dimyristoylated phosphatidylcholine (DMPC) lipids, following established procedures^55,63^. Chloroform-solubilized DMPC (Avanti) was dried under nitrogen gas and left under vacuum overnight to remove residual solvent. The resulting thin film was resuspended in 5% DM (wt vol^-1^) to a final DMPC concentration of 30 mM, and solubilized in a sonicator bath at 37° C. DM-solubilized Cx46/50 (5–6 mg mL^-1^) was combined with DMPC at a molar ratio of 0.6:90 (Cx46/50:DMPC) and incubated at 25° C with gentle agitation for 60 minutes. Purified MSP1E1 was then added at a final molar ratio 0.6:1:90 (Cx46/50:MSP1E1:DMPC) and allowed to incubate at 25° C for an additional 20 minutes. Detergent was removed with SM-2 Bio-Beads (BioRad) at a ratio of 30:1 beads:detergent (wt wt^- 1^) by overnight incubation at 25° C with gentle agitation. Bio-Beads were removed by filtration and the sample was ultra-centrifuged at 146,550 x *g* for 15 minutes at 4° C to remove insoluble material. The supernatant was filtered (Millipore; 0.22 μm) and applied to an SEC column (Superose 6 Increase 10/300 GL; GE Healthcare) equilibrated in 20 mM HEPES (pH 7.4) and 150mM NaCl, to separate empty nanodiscs from Cx46/50-embedded nanodiscs. Peak fractions containing both Cx46/50 and MSP1E1, as determined by SDS-PAGE, were collected and concentrated using a centrifugal device (Vivaspin 6; 50-kDa cut-off filter; Sartorius) to a final concentration ∼2.5 mg mL^-1^, as determined by UV absorbance at 280nm (Extended Data Fig. 1a). All chromatography steps were performed by FPLC at 4° C. The presence of both Cx46 and Cx50 in the final sample was confirmed by western blot analysis using polyclonal antibodies directed against the N-terminal domain of Cx46 (AP11570PU-N, Acris) and the N-terminal domain of Cx50 (LS-C116220, LSBio) (Extended Data Fig. 1b).

### Negative-stain electron microscopy

Cx46/50-lipid nanodisc complexes were prepared for negative stain EM as described^16^. Briefly, a 3 μl drop of sample (∼0.02 mg mL^-1^) was applied to a glow-discharged continuous carbon coated EM specimen grid (Ted Pella), blotted with filter paper and washed two times with detergent-free SEC buffer. The specimen was then stained with freshly prepared 0.75% (wt vol^-1^) uranyl formate (SPI-Chem).

Negatively stained specimens were visualized on a 120kV TEM (iCorr, Thermo Fisher Scientific) at 49,000x magnification at the specimen level (Extended Data Fig. 1c). A total of 76 digital micrographs were collected on a 2k x 2k CCD camera (Eagle 2K TEM CCD, Thermo Fisher Scientific) with a calibrated pixel size of 4.37 Å and with defocus values ranging from 1.5–3.0 μm. All negative-stain image processing was performed in EMAN2.2^64,65^. After contrast transfer function (CTF) parameters were determined, micrographs with significant astigmatism or drift were excluded based on visual inspection of Thon rings in the power spectrum. 7,598 hand-picked particles were extracted with 84 x 84 pixel box size and subjected to multiple rounds of reference-free 2D classification, resulting in a final dataset of 3,826 “good” particles. Representative class averages are shown in (Extended Data Fig. 1c), which revealed dimensions consistent with the expectation that Cx46/50 intercellular channels had been reconstituted into a pair of lipid-nanodiscs.

### CryoEM specimen preparation and data collection

Samples were prepared for CryoEM by applying 5 μl freshly purified Cx46/50-lipid nanodisc complex (∼2.5 mg mL^-1^) to a glow-discharged holey carbon grid (Quantifoil R 1.2/1.3, 400 mesh) for 10 seconds. The grid was blotted for 4.0 seconds and plunge frozen in liquid ethane using a Vitrobot Mark IV (Thermo Fisher Scientific) at 100% humidity and stored under liquid nitrogen.

CryoEM specimen grids were imaged on a Titan Krios (Thermo Fisher Scientific) operated at 300 kV. Dose-fractionated image stacks were recorded on a Falcon 3EC Direct Electron Detector (Thermo Fisher Scientific) at 120,000x nominal magnification in counting mode, with a calibrated pixel size of 0.649 Å pixel^-1^ (Extended Data Fig. 2a). The dose rate was 1.14 e^-^ pixel^-1^ sec^-1^, with 5 frames sec^-1^ collected for a total exposure of 30 seconds, resulting in a total dose for each exposure of ∼52.5 e^-^ Å^-2^. A dataset of 2,087 movies was obtained with nominal defocus values ranging from 1.0–2.2 μm, and data collection parameters were controlled in an automated manner using EPU (Thermo Fisher Scientific).

### Cryo-EM image processing for high-resolution work-flow

The full dataset of 2,087 movies were corrected for beam-induced motion in RELION-3.0^66^ and contrast transfer function (CTF) estimation was performed with Gctf^67^ on the non-dose-weighted, aligned micrographs. Laplacian-of-Gaussian autopicking in RELION-3.0 yielded an initial set of 756,374 picks, which after multiple rounds of 2D classification left 183,784 *bona fide* particles (binned to a 64-pixel box, 3.894 Å pixel^- 1^). These particles were used to generate a *de novo* initial model in RELION, and subsequent 3D refinement of these particles yielded a map at 8.0 Å resolution (64 pixel box, 3.894 Å pixel^-1^). This map was low-pass filtered to 20 Å and projected in 14 unique orientations to perform 3D template-based autopicking in RELION-3.0 to yield 1,210,797 particle picks. Following multiple rounds of 2D classification, this dataset yielded 379,423 “good” particles (200-pixel box, 1.947 Å pixel^-1^) (Extended Data Fig. 2b). Particles that had been translated within 20 Å of their nearest neighbor were removed to prevent invalidation of gold-standard Fourier-shell correlation by duplicate particles. Removal of 120,228 duplicates yielded a 259,195 refined particle set.

This particle set was then re-extracted (1.62 Å pix^-1^, 280-pixel box) and subjected to 3D refinement (D6 symmetry), yielding a map at 3.3 Å resolution. A subsequent round of de-duplication (20 Å cut-off) yielded 227,618 particles that were again re-extracted (0.974 Å pix^-1^, 512-pixel box) and subjected to 3D-refinement (D6 symmetry), which improved the resolution to 3.2 Å. Two rounds of Bayesian polishing and CTF refinement (per-particle defocus, per-micrograph astigmatism) with subsequent 3D refinement (D6 symmetry) yielded a map at 2.7 Å resolution. Particles were then completely unbinned (400-pixel box, 0.649 Å pix^-1^) and subjected to another round of 3D refinement (D6 symmetry), yielding a map that reached the same resolution prior to unbinning (2.7 Å). Bayesian polishing and subsequent 3D refinement of these particles showed no significant improvement.

At this stage, the newly-developed tools in RELION-3.1-beta^68^ were implemented to estimate the degree of beam tilt and high-order aberrations (3-fold and 4-fold astigmatism) present in the particle images. Subsequent 3D refinement (D6 symmetry) improved the resolution to 2.2 Å. Particles that had been translated to within 35 Å of their nearest neighbor (6,224 particles) were again removed to prevent invalidation of the gold-standard Fourier-shell correlation from duplicate particles. The remaining 221,394 particles were subjected to 3D classification into 2 classes with D6 symmetry and a tight solvent mask. Approximately ∼89% of the particles (196,320) fell into one class that was subsequently refined to 2.2 Å resolution (D6 symmetry and solvent mask applied). The remaining 11% of particles (26,005) yielded a 2.0 Å resolution map after 3D refinement (D6 symmetry and solvent mask applied), and all subsequent processing steps were performed on this high-resolution particle set.

Particles were re-extracted with an expanded box size (initially to 448-pixels) to mitigate delocalized CTF signal from particle images with relatively high defocus. New polishing parameters were obtained by running the Bayesian polishing job type in RELION-3.1-beta in “Training mode” on a random 5,000 particle subset of these refined particles. Bayesian polishing was performed with these new parameters and the subsequent 3D refinement (D6 symmetry and solvent mask applied) improved the resolution slightly to 1.97 Å. This process was iterated multiple times with successive increase in box size and incrementally tighter solvent mask applied during Bayesian polishing until no further improvements were observed, resulting in a final box size of 540 pixels and refined map at 1.94 Å resolution with D6-symmetry and 2.3 Å resolution without symmetry (Gold-Standard, 0.143 cut-off)^69^ (Extended Data Fig. 2c, 3a). Local resolution of the final map was estimated in RELION-3.1-beta^68^, and local resolution-filtered maps were generated for model building (Extended Data Fig. 2d, 3b). A schematic illustrating this high-resolution CryoEM workflow is presented in (Extended Data Fig. 2c).

### Cryo-EM image processing workflow for lipid classification

For classification and analysis of lipid configurational/conformational heterogeneity, a modified workflow starting from the totally unbinned 227,618 particle set (0.649 Å pix^-1^, 400-pixel box) which yielded the 2.7 Å resolution map was applied, as described here (and illustrated in Extended Data Fig. 4a). The particle set was subjected to 3D classification (eight classes), with D6 symmetry and a generous solvent mask applied. Two of the eight classes yielded maps in which the lipid configuration was unambiguously resolved: assigned as PC Class 1, containing 9,190 particles (∼4% of the data) and PC Class 3, containing 6,944 particles (∼3% of the data). Overlapping configurations were resolved in two of the other 3D classes, and so particles from these classes were combined and subjected to a second round of 3D classification with only 2 classes and a tight solvent mask applied. This yielded one class with unresolved lipid configurations, and a second class in which the lipid configuration was unambiguously resolved: assigned PC Class 2, containing 6,075 particles (∼3% of the data). Particles assigned to PC Class 1, 2, and 3 were separately subjected to a final round of 3D refinement with a solvent mask and D6 symmetry applied (Extended Data Fig. 4a). The final reconstructions from particles in each of these classes all reached ∼2.5 Å resolution (Gold-Standard, 0.143 cut-off) (Extended Data Fig. 4b). Local resolution was estimated in RELION-3.1-beta, and local resolution-filtered maps were generated for model building (Extended Data Fig. 4c).

### Atomic modelling, refinement and validation

For all atomic models of Cx46 and Cx50, initial models were derived from previously reported CryoEM structure of amphipol-stabilized Cx46 and Cx50 (PDB 6MHQ and 6MHY^16^, respectively). Initial models were fit as rigid bodies into the D6-symmetrized CryoEM maps with applied local resolution-filtering using UCSF Chimera^70^. All atom models for Cx46 and Cx50 were further built into the CryoEM density maps with COOT^71^, and subjected to real-space refinement in PHENIX^72^ with secondary structure and non-crystallographic symmetry (D6) restraints applied. Several iterations of manual adjustment of the protein model in COOT followed by real-space refinement in PHENIX, were performed while monitoring model quality with MolProbity^73^ and quality of side chain fit with EMRinger^74^. Coordinate and restraint files for the dimyristoylated phosphatidylcholine (DMPC) ligands were generated with PHENIX eLBOW^75^. DMPC molecules were manually fit into the Cryo-EM density with COOT. Since density for the phosphatidylcholine (PC) head groups was not resolved in the high-resolution ensemble CryoEM map (1.9 Å map), head group and acyl chain atoms that could not be accommodated by the density were deleted. For the PC Lipid Classes 1–3, the postprocessed maps from RELION were low pass-filtered to 3.5 Å resolution to facilitate modeling of the fully-resolved PC lipids. COOT was further used to manually place water molecules into solvent densities of the CryoEM maps. Appropriate placement of waters was determined by the following three criteria: 1) confirmation of at least two hydrogen bond donor/acceptor interactions with the FindHBond tool in UCSF Chimera (< 4 Å donor-acceptor distance), 2) confirmation of solvent densities consistently observed in both gold-standard separated half-maps (contoured ≥ 2.5 σ), and 3) as an additional measure we looked for density overlap between the local resolution-filtered Cryo-EM map (contoured ≥ 5.3 σ) and the time-averaged water density map generated by equilibrium molecular dynamics simulation (contoured ≥ 5.0 σ) to help assign weak experimental water densities (Extended Data Fig. 8, see calculation of water density maps from MD described in Methods below). However, not all of the assigned CryoEM water densities were observed by MD (76% of waters were observed at equivalent positions by CryoEM and MD). Several iterations of real-space refinement on the entire model were completed until refinement statistics converged.

### Disclosure of unresolved heteromeric/heterotypic assemblies of Cx46/50

All models of Cx46 and Cx50 were built using D6 (12-fold) symmetrized CryoEM maps. Because native Cx46/50 intercellular channels may form homomeric and/or various patterns of heteromeric/heterotypic configurations^16,76,77^, this map most likely represent a heterogeneous mixture of these two isoforms^16^. This approach was chosen because all attempts to separate the heteromeric/heterotypic assembly of these two isoforms using image classification procedures were unsuccessful (presumably due to the close sequence and structural similarity of these two isoforms) (see also^16^). Indeed, Cx46 and Cx50 are 80% identical and 89% similar in sequence over the resolved structural domains, while sites of difference are typically at solvent exposed regions (Extended Data Fig. 6a). Despite this limitation, all atomic-models generated by this approach showed good stereo-chemical refinement statistics (see Extended Data Table 1), and significant improvements to the previously described amphipol-stabilized models that were refined to 3.4 Å resolution (Extended Data Fig. 5a-e). It is important to note that sites in the density maps where the sequence of Cx46 and Cx50 are identical or similar, both models fit well into the D6 symmetrized map, and these regions tend to display well-resolved sidechain density (Extended Data Fig. 3c,d). Over regions where the sequence of Cx46 and Cx50 differ, sidechain density is sometimes weaker and/or displays appearance of density consistent with a mixture of both isoforms (Extended Data Fig. 3c,d). These observations are possibly due to the imposed D6 symmetry averaging of density belonging to two different sidechains in these areas, or relative flexibility at these sites as many of these residues are also solvent-exposed. In these areas of difference, where EM density is observed, both Cx46 and Cx50 can be fit into the density equally well (Extended Data Fig. 3c,d). Nevertheless, caution should be used with interpretation of the conformational details at these sites of isoform difference.

### Molecular dynamics simulations

Visual Molecular Dynamics (VMD) v1.9.3^78^ was used to build systems for sheep Cx46 and Cx50 in a dual lipid-bilayer with varying salt conditions, designed to mimic either the cellular environment (cytoplasmic KCl, extracellular NaCl) or experimental CryoEM conditions (uniform NaCl). To produce unbiased analysis of water and lipid interactions, all water and lipid molecules derived by CryoEM analysis were removed from the Cx46 and Cx50 models prior to the MD setup. Each system comprised the full dodecameric gap junction intercellular channel, prepared in explicit water (model TIP3P) and embedded in two lipid bilayers composed of dimyristoylated phosphatidylcholine (DMPC), mimicking the cell-to-cell junction. For all models, sidechains were protonated according to neutral conditions and the HSD model was used for all histidine residues. Disulfide bonds identified in the experimental structures were enforced. Amino acids corresponding to the intracellular loop (ICL; residues 110–136 in sheep Cx46 and residues 110–148 in sheep Cx50) and c-terminal domain (CTD; residues 225–413 in sheep Cx46 and residues 237–440 in sheep Cx50) were not included for the MD simulations, as experimental data describing the structure of these large flexible domains (∼30 residue ICL and ∼200 residue CTD in Cx46 and Cx50) are missing. The introduced n- and c-terminal residues resulting from the missing ICL segment (sheep Cx46 R109 and K137; sheep Cx50 R109 and R149) were neutralized. All of the systems were modified with an n-terminal acetylation (at the starting residue Gly 2) in VMD through an all-atom acetylation patch in the automated PSF-Builder, in accordance with previously described proteomics analysis on native Cx46/50^16,59,79^, and expectation that this species would predominate in cells^80^. A complete list of modeled residues for each system is provided in Extended Data Fig. 7a.

The prepared protein structures were submerged in a hydration shell using Solvate 1.0.1^81^. Water was removed from sections of the channel corresponding to transmembrane domains, based on hydrophobic character and localization of lipid-nanodisc observed in the experimental CryoEM data (+/- 20–50 Å from the center of the channel). The CHARMM-GUI membrane-builder^82^ was used to build the DMPC bilayers (pre-melted), with dimensions of 154 x 154 Å for Cx46 and Cx50, and lipids overlapping with protein were removed. The entire system was then placed in a water box with dimensions 147 x 147 x 174 Å for both Cx46 and Cx50, using VMD’s Solvate plugin. The system was neutralized using the Autoionize plugin, then 150 mM KCl and 150 mM NaCl was added to the solvent areas corresponding to intracellular and extracellular regions of the simulation box for the “KCl” systems, while the “NaCl” systems contained 150 mM NaCl for the entire box. A summary of atoms counts for each system is provided in Extended Data Fig. 7a.

CUDA-accelerated nanoscale molecular dynamics (NAMD) 2.13^83^ was used for all classical MD simulations, using the CHARMM36 force-field^84^ for all atoms and TIP3P explicit model for water. Each system was prepared following the same minimization and equilibration protocol, as follows. An initial minimization step, where the lipids, solvent and ions were allowed to minimize around the protein was performed, with the protein harmonically constrained for 1 ns, with 1 fs timestep and constant pressure (NPT ensemble). A second minimization step was applied, where the system was free to minimize with a harmonic constraint on the protein backbone to ensure stable quaternary structure for 1 ns – lipids relax and compress during minimization steps with minimized dimensions equal to the water box (14.7 x 14.7). The entire system was then released from restraints and subjected to all-atom equilibration runs employing Langevin thermostat, with a constant temperature of 310 K and constant pressure of 1 atm (NPT ensemble), with 2 fs time-steps and allowed to proceed for 30 ns. Periodic boundary conditions were used to allow for the smoothed particle mesh Ewald (PME) calculation of electrostatics. Finally, two independent 100 ns production runs were seeded with randomly initialized velocities from the initial equilibration simulation – providing 200ns of production for each system. Root mean squared deviations (r.m.s.d.) and root mean squared fluctuations (r.m.s.f.) were calculated using VMD, and r.m.s.f. values were displayed to the protein structure using UCSF Chimera (Extended Data Fig. 7b-d). All systems approached a steady r.m.s.d. within 30 ns of the equilibration phase (Extended Data Fig. 7b), and r.m.s.f. values appeared well-behaved over the production periods, including regions corresponding to the NTH domain^16^ (Extended Data Fig. 7c,d). The only significant fluctuations (i.e., > 2.5 Å) occurred at the TM2, TM3 and TM4 cytoplasmic termini, which is expected as these regions form the boundary to the intrinsically disordered ICL and CTD regions of the protein (not modeled). All systems maintained an electro-chemical seal to extracellular sodium ions (Na^+^) around the ECD docking domains during MD simulation.

### Calculation of density maps from MD for water, lipids and ions

The Volmap plugin in VMD was used for the calculation of volumetric density maps, by replacing each atom with a normalized gaussian distribution, whose standard deviation is equal to the radius of the atom. All of the gaussians are summed and distributed on a grid for each frame of the simulation. The grids were re-sampled to a final voxel resolution of 0.649 Å to match the pixel size used in the CryoEM reconstruction. Water, ion, and lipid maps were calculated from each of two 100 ns production runs, and subsequently averaged and symmetrized (D6-symmetry) with the relion_image_handler tool in RELION-3.0^66^. Lipid and water density maps produced from Cx46 and Cx50 MD simulations contained significant overlap to each other and to the CryoEM maps, however, the maps produced from Cx50 MD simulations were of higher quality and were selected for detailed comparative analysis to the CryoEM density maps (Fig. 3 and Extended Data Fig. 8). Ion density maps showed only a few features and did not correspond to densities observed by CryoEM (not shown), and were therefore excluded from further analysis.

### MD-based area per lipid (APL) and lipid order parameter (S_CD_) calculations

Area per lipid (APL) for each membrane, separated by intracellular and extracellular leaflet, were calculated using the program FATSLiM^85^, and used as an indicator of equilibration of the lipid systems (Extended Data Fig. 9a).

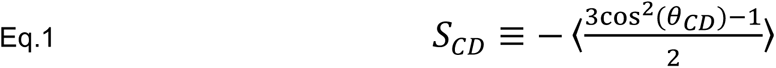

The S_CD_ lipid order parameter, as defined by Eq. 1, measures the orientation of the SN1 and SN2 acyl-chains by monitoring the angle that each acyl C-H vector makes with the bilayer normal *θ*_*CD*_. The calculations of S_CD_ were done using the VMD script *calc_op*.*tcl*^86^. To analyze the distance dependence of S_CD_ in the respective membrane leaflets, the averaged S_CD_ values were calculated in 5 Å concentric shells around the protein (SN1 and SN2 calculated separately). S_CD_ of lipids from both membranes are averaged together, while the intra- and extracellular leaflets were averaged separately (Extended Data Fig. 9b-e). To visualize the order parameter mapped to the structure, the time-averaged S_CD_ values were calculated for each lipid (SN1 and SN2 combined values for acyl carbons 4–11), and colored according to this value using UCSF Chimera (Fig. 3c).

### MD lipid configuration analysis

Analysis of PC lipid configuration (*i*.*e*., acyl-chain positioning) was performed using in-house scripts to assess how phospholipids are organized within the extracellular leaflet of the Cx46 and Cx50 intercellular channels during MD simulation, as compared to the PC configurations classified by CryoEM. This was done by counting the instances of a single DMPC molecule occupying the region bounded by the MD-based lipid-density, contoured at σ_*min*_ = 8 (Fig. 3a). The lipid acyl-chain density maps calculated from the Cx50 MD simulations reveal more than 19 resolved rods (*i*) of density per connexin subunit (12 subunits), and each rod was arbitrarily numbered 1 through 19 (total of 228 acyl-chain positions). A lipid was classified in a state when both acyl-chains occupied a density, *state* ≡ “*i* − *j*” (*where i* ≠ *j*). A rod density is considered occupied if at least 5 carbons of a lipid’s acyl-chain are within the density, such that σ_*carbon*_ ≥ σ_*min*_. This classification scheme was applied to every lipid within 15 Å of the protein, over each frame (0.1 ns per frame). To analyze the dynamics of lipids surrounding the protein, the state of each lipid (e.g., “1–2”, “1–5”, “none”, etc.) was monitored and recorded at every frame providing a time series of lipid configurational dynamics in state-space (Fig. 4d).

## Statistical analysis

95% confidence intervals for Cα r.m.s.f. values are reported (n=24) using a two-tailed student t-test. Fourier-Shell Correlation (FSC) was performed using Gold-Standard methods with a 0.143 cut-off criteria^69^. No statistical methods were used to predetermine sample size for the CryoEM dataset. The experiments were not randomized, and investigators were not blinded to allocation during experiments and outcome assessment.

## Data Availability

CryoEM density maps have been deposited to the Electron Microscopy Data Bank (EMD-XXXX). Coordinates for Cx46 and Cx50 atomic models have been deposited to the Protein Data Bank (XXXX and XXXX). The original multi-frame micrographs have been deposited to EMPIAR (EMPIAR-XXXXX).

## Extended Data Tables and Figures

**Extended Data Table 1.** CryoEM Statistics. Summary of CryoEM data collection, refinement and model validation statistics. The ensemble CryoEM dataset was used to obtain the 1.9 Å resolution reconstruction and atomic models for Cx46 and Cx50, including 396 water molecules and 150 lipid acyl-chains. 3D classification was used to obtain the three PC classes, and associated atomic models for Cx46 and Cx50 (PC Class 1–3). Pre-processed and post-processed maps and associated masks from all datasets have been deposited to the EM databank (EMD-XXXX). The original multi-frame micrographs have been deposited to EMPIAR (EMPIAR-XXXX). Coordinates for Cx50 and Cx46 atomic models have been deposited to the Protein Data Bank (XXXX and XXXX correspond to the ∼1.9 Å models, XXXX and XXXX correspond to the ∼2.5 Å models from PC Class 1; XXXX and XXXX correspond to PC Class 2; and XXXX and XXXX correspond to PC Class 3).

|  | Cx46 | Cx50 | Cx46<br>*PC Class 1* | Cx50<br>*PC Class 1* | Cx46<br>*PC Class 2* | Cx50<br>*PC Class 2* | Cx46<br>*PC Class 3* | Cx50<br>*PC Class 3* |
| --- | --- | --- | --- | --- | --- | --- | --- | --- |
| **Data collection and processing** |  |  |  |  |  |  |  |  |
| Magnification | 120,000 | 120,000 | 120,000 | 120,000 | 120,000 | 120,000 | 120,000 | 120,000 |
| Voltage (kV) | 300 | 300 | 300 | 300 | 300 | 300 | 300 | 300 |
| Electron exposure (e-/Å<sup>2</sup>) | 52.5 | 52.5 | 52.5 | 52.5 | 52.5 | 52.5 | 52.5 | 52.5 |
| Defocus range (μm) | -1.0 to -2.2 | -1.0 to -2.2 | -1.0 to -2.2 | -1.0 to -2.2 | -1.0 to -2.2 | -1.0 to -2.2 | -1.0 to -2.2 | -1.0 to -2.2 |
| Pixel size (Å) | 0.649 | 0.649 | 0.649 | 0.649 | 0.649 | 0.649 | 0.649 | 0.649 |
| Symmetry imposed | D6 | D6 | D6 | D6 | D6 | D6 | D6 | D6 |
| Initial particle images (no.) | 1,210,797 | 1,210,797 | 1,210,797 | 1,210,797 | 1,210,797 | 1,210,797 | 1,210,797 | 1,210,797 |
| Final particle images (no.) | 26,005 | 26,005 | 6,073 | 6,073 | 9,188 | 9,188 | 6,942 | 6,942 |
| Map resolution (Å) | 1.94 | 1.94 | 2.52 | 2.52 | 2.47 | 2.47 | 2.45 | 2.45 |
| FSC threshold | 0.143 | 0.143 | 0.143 | 0.143 | 0.143 | 0.143 | 0.143 | 0.143 |
| Map resolution range (Å) | 1.94–3.30 | 1.94–3.30 | 2.42–4.56 | 2.42–4.56 | 2.33–4.14 | 2.33–4.14 | 2.47–4.33 | 2.47–4.33 |
| **Refinement** |  |  |  |  |  |  |  |  |
| Initial model used (PDB code) | 6MHQ | 6MHY | 6MHQ | 6MHY | 6MHQ | 6MHY | 6MHQ | 6MHY |
| Model resolution (Å) | 1.9 | 1.9 | 2.5 | 2.5 | 2.4 | 2.4 | 2.6 | 2.7 |
| FSC threshold | 0.5 | 0.5 | 0.5 | 0.5 | 0.5 | 0.5 | 0.5 | 0.5 |
| Map sharpening *B* factor (Å<sup>2</sup>) | -26.28 | -26.28 | -40.49 | -40.49 | -45.42 | -45.42 | -43.73 | -43.73 |
| **Model composition** |  |  |  |  |  |  |  |  |
| Non-hydrogen atoms | 21,876 | 21,804 | 22,308 | 22,176 | 21,624 | 21,552 | 21,312 | 21,240 |
| Protein residues | 2352 | 2352 | 2,328 | 2,328 | 2,316 | 2,316 | 2,304 | 2,304 |
| Ligands | 180 | 180 | 168 | 168 | 132 | 132 | 144 | 144 |
| ***B* factors (Å<sup>2</sup>)** |  |  |  |  |  |  |  |  |
| Protein | 31.48 | 30.10 | 51.09 | 50.04 | 41.92 | 47.13 | 59.04 | 57.75 |
| Ligand | 32.95 | 30.94 | 55.25 | 55.19 | 43.92 | 47.64 | 61.77 | 60.83 |
| **R.m.s. deviations** |  |  |  |  |  |  |  |  |
| Bond lengths (Å) | 0.006 | 0.006 | 0.010 | 0.007 | 0.009 | 0.009 | 0.007 | 0.010 |
| Bond angles (°) | 0.784 | 0.785 | 0.825 | 0.821 | 0.934 | 1.073 | 0.816 | 0.762 |
| **Validation** |  |  |  |  |  |  |  |  |
| MolProbity score | 1.09 | 1.07 | 1.03 | 1.03 | 1.13 | 1.22 | 0.98 | 1.06 |
| Clashscore | 2.98 | 2.61 | 2.45 | 2.46 | 2.90 | 3.93 | 2.07 | 2.72 |
| Poor rotamers (%) | 0.57 | 0.56 | 0.00 | 0.56 | 1.17 | 1.14 | 0.00 | 0.57 |
| **Ramachandran plot** |  |  |  |  |  |  |  |  |
| Favored (%) | 98.44 | 97.92 | 98.11 | 98.95 | 98.41 | 98.94 | 98.40 | 98.40 |
| Allowed (%) | 1.56 | 2.08 | 1.89 | 1.05 | 1.59 | 1.06 | 1.60 | 1.60 |
| Disallowed (%) | 0.00 | 0.00 | 0.00 | 0.00 | 0.00 | 0.00 | 0.00 | 0.00 |

**Extended Data Figure 1.**
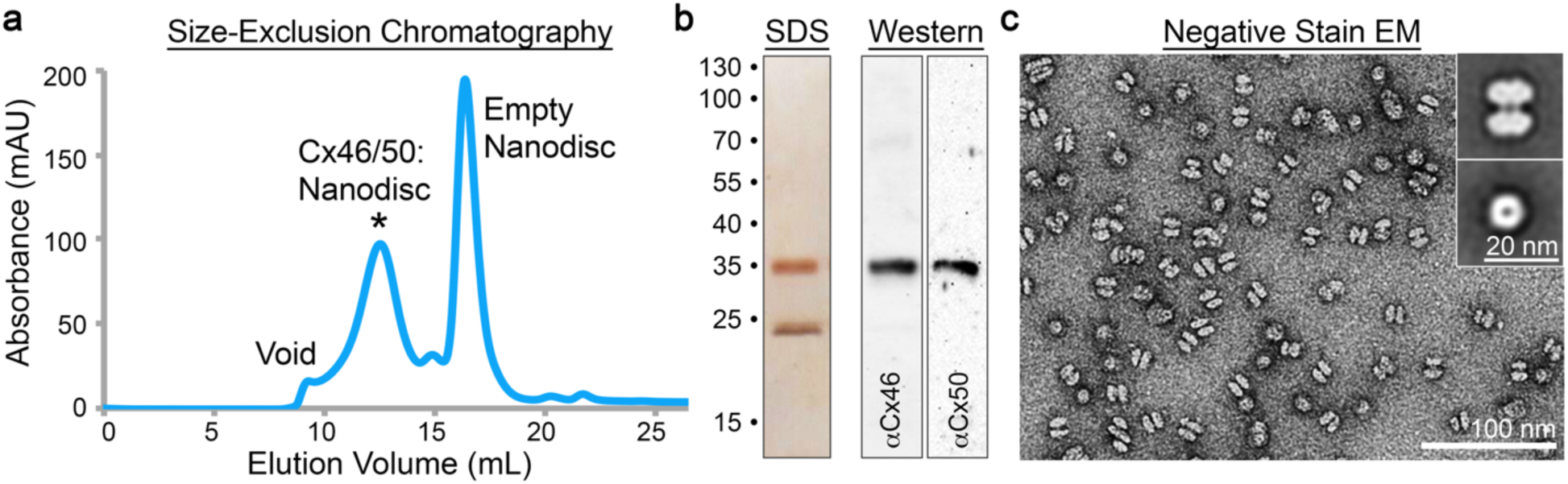
Cx46/50 reconstitution into MSP1E1/DMPC lipid-nanodiscs and negative stain EM. **a)** Size-exclusion chromatography (SEC) trace monitored by UV absorbance at 280 nm. Peaks corresponding to Cx46/50 reconstituted into MSP1E1/DMPC nanodiscs (*), empty nanodisc and void are indicated. **b)** SDS-PAGE (*left*) and western blot (*right*) of peak SEC fraction (labeled *), with molecular weight markers indicated. MSP1E1 migrates as a ∼24 kDa band (predicted MW ∼27.5 kDa). Cx46 and Cx50 migrate together at ∼38 kDa band, as expected from c-terminal truncation from core lens fiber cells^16^, and confirmed by western blot (*right*). **c)** Electron micrograph of negatively stained particles from SEC fraction (labeled *), with scale bar = 100 nm. *Inset*, shows representative 2D class averages of sideview (*top*) and top view (*bottom*), with scale bar = 20 nm. Particles display dumbbell-like structures corresponding to Cx46/50 gap junctions intercellular channels^16,87^, embedded into a pair of ∼10-11 nm wide nanodiscs (MSP1E1 nanodiscs have a predicted diameter of ∼10.5 nm^17^).

**Extended Data Figure 2.**
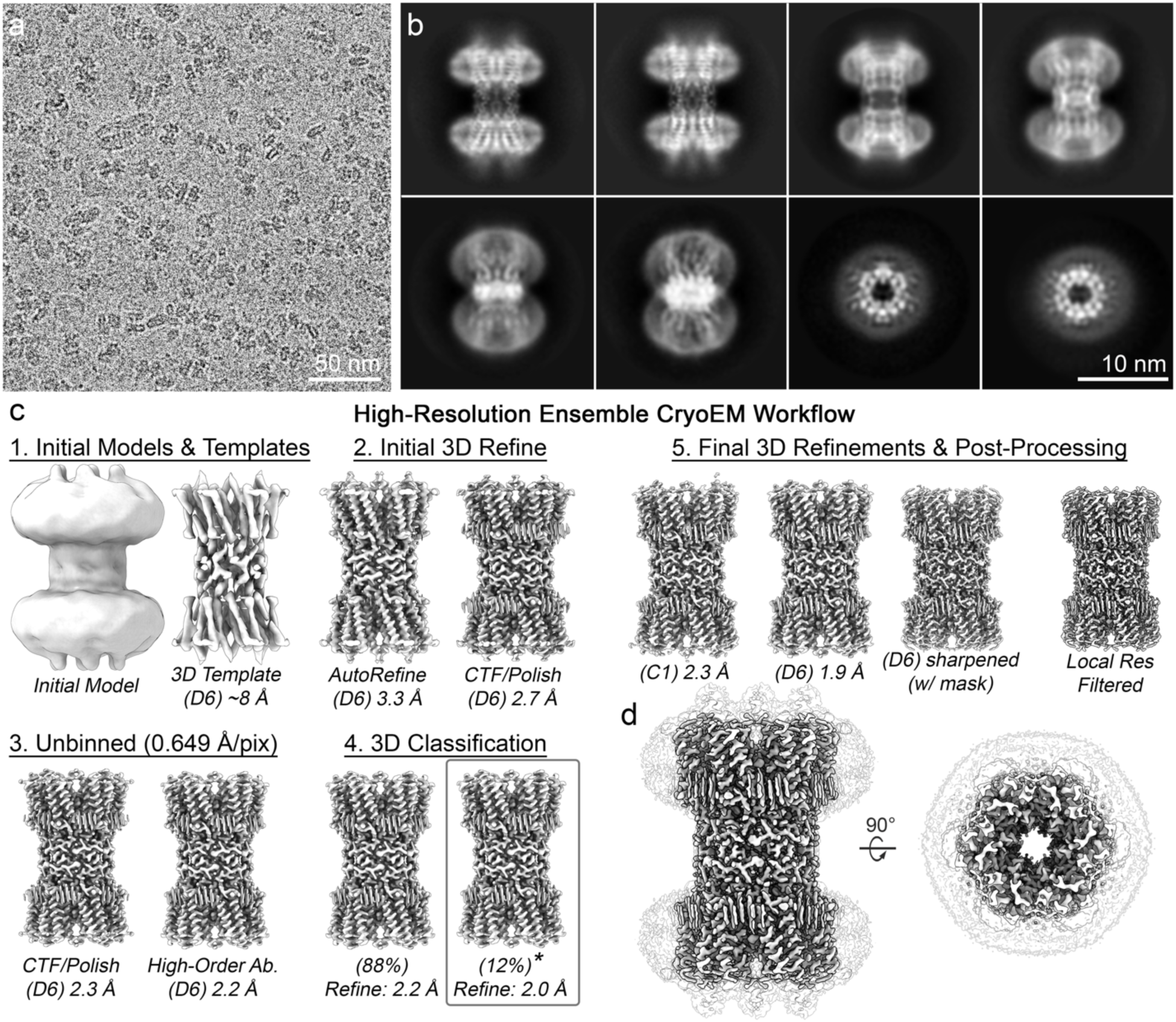
CryoEM image processing workflow for 1.9 Å ensemble reconstruction of Cx46/50 in DMPC lipid nanodiscs. **a)** Representative CryoEM micrograph (dataset of 2088 movies) recorded on a Falcon III detector, with physical pixel size = 0.649 Å^2^ and total dose of ∼60 e^-^ per Å^2^. Scale bar = 50 nm. **b)** Representative 2D class averages. Scale bar = 10 nm. **c)** Image processing and 3D reconstruction workflow carried out in Relion^66,68^, with representative maps at different stages of the image processing pipeline. Step 1) *De-novo* model generated in Relion (*left*) and initial 3D AutoRefinement with D6-symmetry (∼8 Å resolution, 3.9 Å pixel size) (*right*), which was then filtered to 20 Å and used for 3D template auto-picking in Relion (resulting in ∼1.2M particle picks, which were culled to ∼228k “good” particles following multiple rounds of 2D classification and de-duplication). Step 2) Resulting 3D AutoRefine with D6-symmetry (3.2 Å resolution, 0.97 Å pixel size) (*left*), and resulting map following per particle CTF-refinement and polishing in Relion (2.7 Å) (*right*). 3) Particles were unbinned (pixel size 0.649 Å/pix, box size = 400 pix) and refined with per-particle CTF-correction and polishing (2.3 Å) (*left*), and further refinement of high-order aberration parameters in Relion v3.1-beta^68^ (2.2 Å) (*right*). Step 3) Particles were de-duplicated, resulting in a set of ∼221k particles, and subjected to 3D classification (two classes). Class 1 contained 88% of the particles and was further refined to 2.2 Å resolution (*left*). Class 2 contained 12% of the particles and was further refined to 2.0 Å resolution (*right, asterisk*). Step 4) Particles belonging to Class 2 (∼26k particles), were then subjected to multiple rounds of 3D Auto-refinement followed by per-particle CTF, aberration-correction and polishing, using successively larger box-sizes until no further improvement, resulting in a final reconstruction at 2.3 Å resolution (C1 symmetry) (*left*) and 1.9 Å resolution (D6 symmetry) (*center, left*). The D6-symmeterized map was then subjected to post-processing (b-factor sharpening) (*center, right*) and local-resolution filtering in Relion (*right*) for downstream analysis. **d)** Final reconstruction of Cx46/50 following local resolution filtering, used for atomic-modeling. Transparent silhouette displays the unmasked map at low-contour to illustrate the dimensions of the lipid nanodisc densities.

**Extended Data Figure 3.**
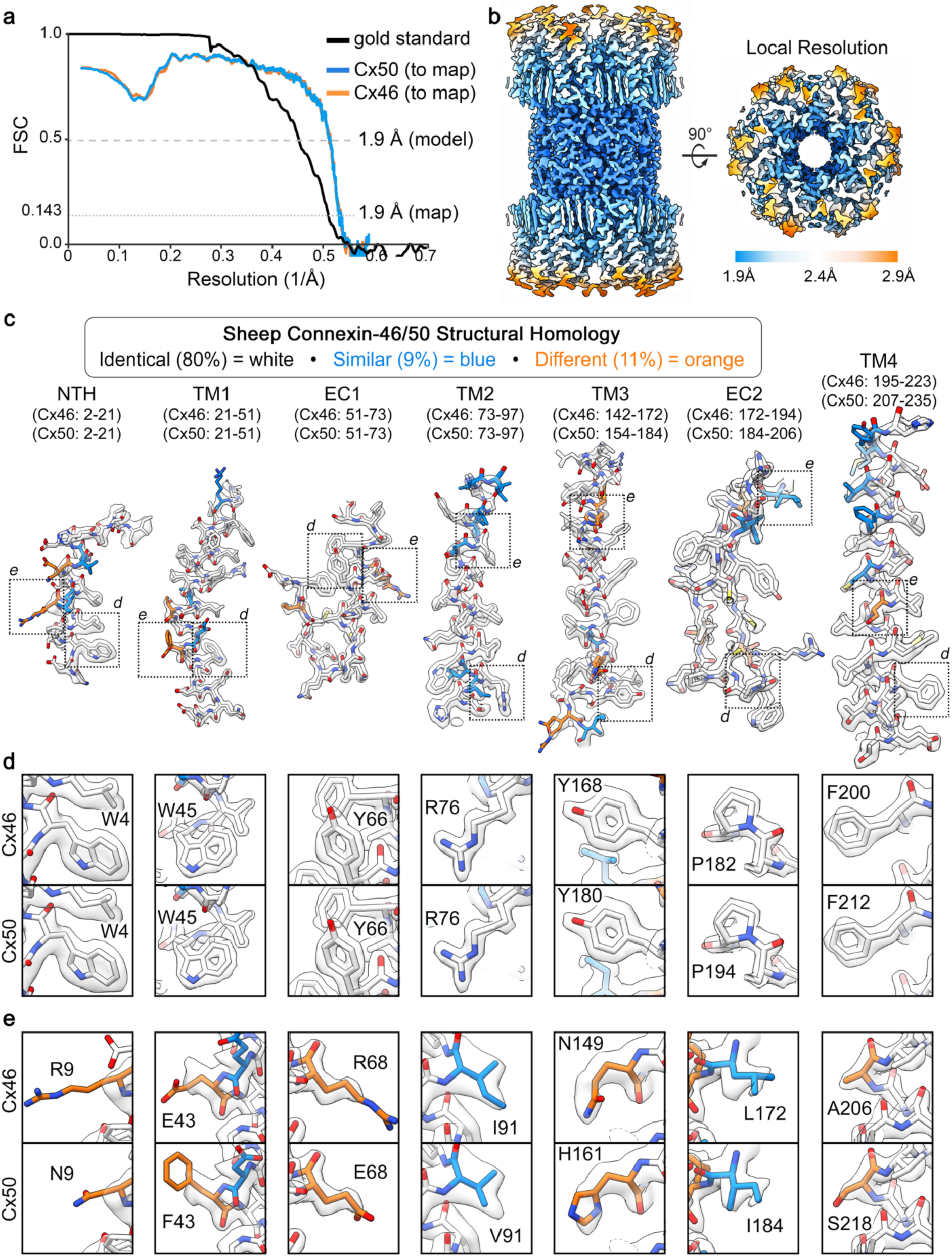
Global and local resolution assessment of the 1.9 Å ensemble reconstruction of Cx46/50 in DMPC lipid nanodiscs. **a)** Fourier Shell Correlation (FSC) analysis obtained from the ensemble CryoEM map of Cx46/50 in DMPC lipid nanodiscs. Gold-standard FSC (black) of the final refined CryoEM map indicates a global resolution of 1.9 Å (0.143 cut-off). FSC curves comparing atomic models for Cx50 (blue) and Cx46 (orange) fit to the final CryoEM map display overall correlation at 1.9 Å (0.5 cut-off). **b)** Local resolution analysis of the final CryoEM map using Relion^68^, displayed by colored surface (1.9 – 2.4 Å = blue – white; 2.4 – 2.9 Å = white – orange). **c)** Segmented CryoEM map with regions of the atomic models for sheep Connexin-46 (Cx46) and Connexin-50 (Cx50) fit to the local-resolution filtered map. Residue numbering for Cx46 and Cx50 is displayed above the corresponding segments for the n-terminal helix domain (NTH) domain, the transmembrane domains 1-4 (TM1-4) and extracellular domains 1-2 (EC1-2), Residues are colored according to the pair-wise sequence homology between sheep Cx46 and Cx50, as being identical (white, 80%), similar (blue, 9%) and different (orange, 11%), with all heteroatoms colored by standard scheme (oxygen – red; nitrogen – blue, sulfur – yellow). **d, e)** Windows show zoom-views corresponding to boxed regions of the segmented maps. **d)** Displays fits over representative regions where both Cx46/50 contain identical amino acids, where the high-resolution features are well-resolved. **e)** Displays fits over representative regions where the sequence of Cx46 and Cx50 differ, and where sidechain density is weaker and/or consistent with heterogeneity. This is presumably due to the heteromeric/heterotypic mixture of these isoforms^16,76,77^ and the imposed averaging of two different sidechains in these areas, and/or to relative flexibility at these sites, as many of these same residues correspond to solvent/lipid exposed sidechains (e.g., R9/N9; E43/F43, R68/E68, I91/V91 and A206/S218).

**Extended Data Figure 4.**
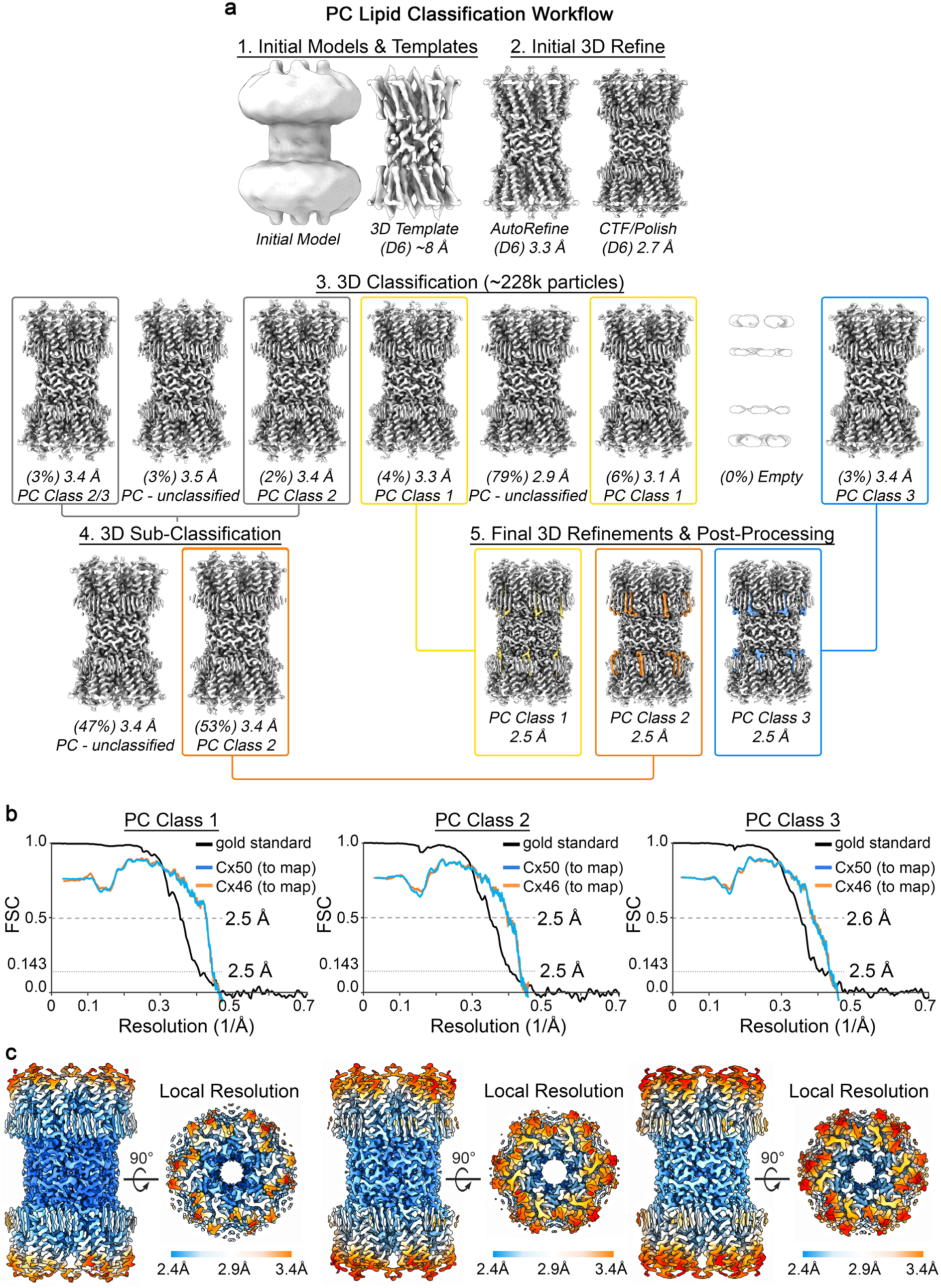
Image processing and resolution assessment for 3D lipid-classification work-flow. **a)** Image process and 3D reconstruction workflow carried out in Relion for the analysis of PC lipid configuration/conformational heterogeneity, with representative maps at different stages of the image processing pipeline. Steps 1 and 2) are the same as described in Extended Data Fig. 2, which resulted in a 2.7 Å reconstruction from a dataset of ∼228k “good” particles (*right*). Step 3) These particles were unbinned and re-extracted (0.649 Å/pix, 400 pixel box), and subjected to 3D classification (eight classes) without image alignment. Two of the eight classes yielded maps in which the lipid configuration was unambiguously resolved: assigned as PC Class 1 (yellow box), containing 9,190 particles (∼4% of the data) and PC Class 3 (blue *box*), containing 6,944 particles (∼3% of the data). Overlapping configurations were resolved in two of the other 3D classes (grey boxes). Step 4) The particles from these overlapping classes (grey boxes) were combined and subjected to a second round of 3D classification with two classes. This yielded one in which the lipid configuration was unambiguously resolved: assigned PC Class 2 (orange box), containing 6,075 particles (∼3% of the data). Step 5) Particles assigned to PC Class 1 (*left*), PC Class 2 (*center*) and PC Class 3 (*right*) were separately subjected to a final round of 3D refinement and per-particle polishing, with D6 symmetry applied, resulting in final reconstructions ∼2.5 Å resolution (Gold-Standard, 0.143 cut-off). **b)** Fourier Shell Correlation (FSC) analysis obtained for PC Class 1 (*left*), PC Class 2 (*center*) and PC Class 3 (*right*). Gold-standard FSC (black) of the final refined, masked and post-processed CryoEM map indicates a global resolution of 2.5 Å (0.143 cut-off). FSC curves comparing atomic models for Cx50 (blue) and Cx46 (orange) fit to the final CryoEM maps display overall correlation at 2.5–2.6 Å (0.5 cut-off). **c)** Local resolution analysis of the final CryoEM maps for PC Class 1 (*left*), PC Class 2 (*center*) and PC Class 3 (*right*) using Relion, displayed by colored surface (2.4 – 2.9 Å = blue – white; 2.9 – 3.4 Å = white – orange).

**Extended Data Figure 5.**
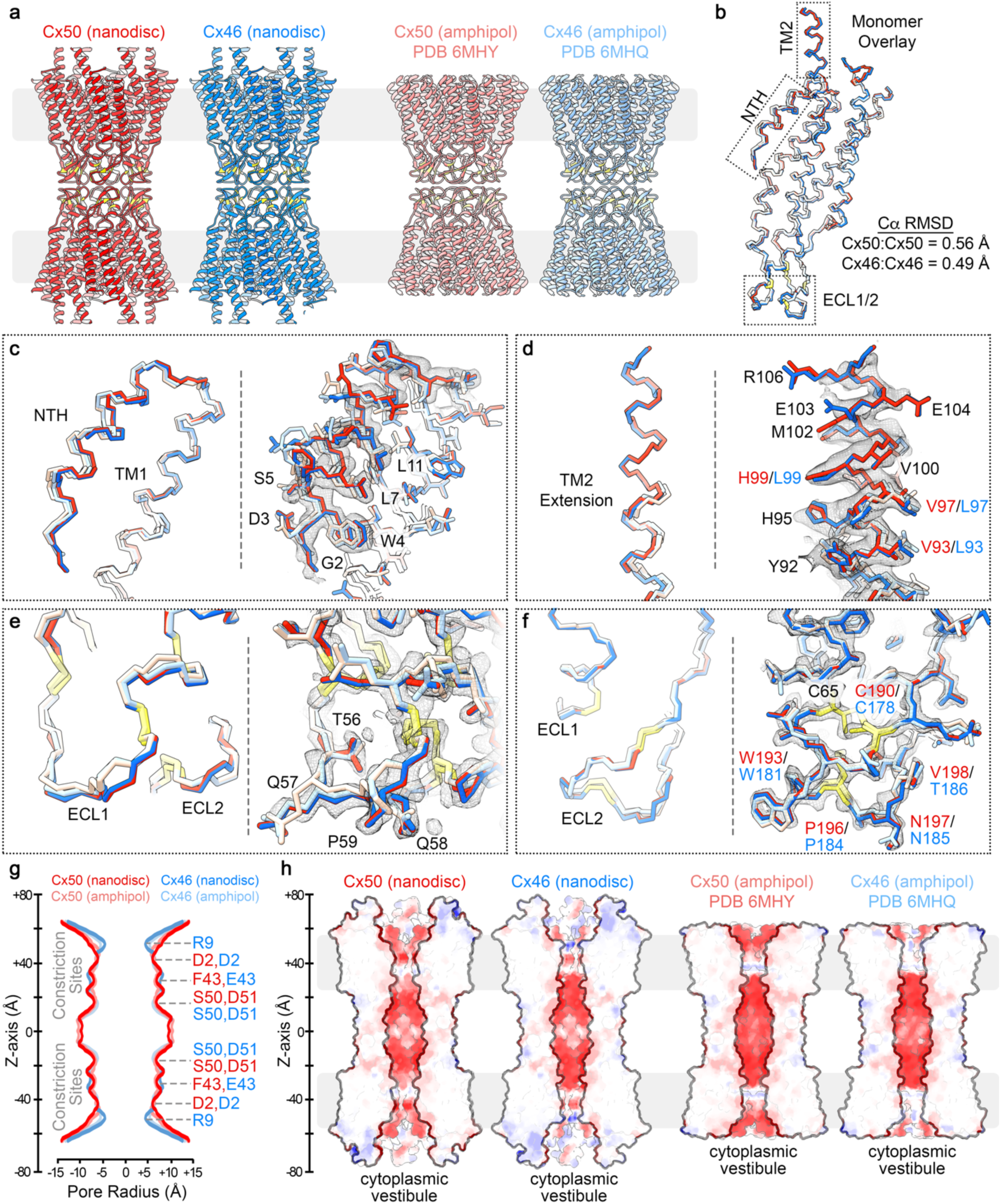
Comparison of Cx46/50 structures determined in amphipol and lipid-nanodiscs. **a)** Ribbon structures of Cx50 (red) and Cx46 (blue) determined by CryoEM in lipid-nanodisc (left) and as previously determined in amphipol (right) with Cx50 (light red, PDB 6MHY) and Cx46 (light blue, PDB 6MHQ)^16^. Regions of lipid bilayer are indicated by light grey box. Conserved cysteine positions within the EC1/2 domains, involved in disulfide formation, are indicated in yellow. **b)** Cα traces over-laid for these four models, corresponding to a single subunit following super-positing (colored as in panel a). Cα r.m.s.d. following super-positioning is indicated for Cx50 (nanodisc) vs. Cx50 (amphipol) = 0.56 Å, and Cx46 (nanodisc) vs. Cx46 (amphipol) = 0.49 Å. **c–f)** Shows zoom views corresponding to the boxed regions in panel b. For each panel, (left) shows Cα trace and (right) shows all atom fit into the 1.9 Å CryoEM density map obtained from the nanodisc embedded structure, to show regions of improved fit to the experimental density map. Highlighted residues are indicated, and labels colored according to identity between the Cx50 and Cx46 isoforms (black – identical, red – Cx50 and blue – Cx46). **g)** Pore radius determined using HOLE^88^, for experimental structures of Cx50-nanodisc (red), Cx46-nanodisc (blue), Cx50-amphipol (light red) and Cx46-amphipol (light blue). Locations corresponding to constriction sites are indicated, and residues contributing to these sites of constriction for both isoforms are labeled (Cx50 – red; Cx46 – blue). **h)** Cut-away surface representation of Cx50-nanodisc *(left)*, Cx46-nanodisc *(left center)* and Cx50-amphipol *(right center)* and Cx46-amphipol *(right)*, colored by coulombic potential (negative – red, neutral – white and positive – blue). This comparison illustrates the electrostatic environment of the permeation pathways and the extension of the intra-cellular vestibule that is resolved in the Cx46/50-nanodisc models, as compared to the previously described Cx46/50-amphipol models.

**Extended Data Figure 6.**
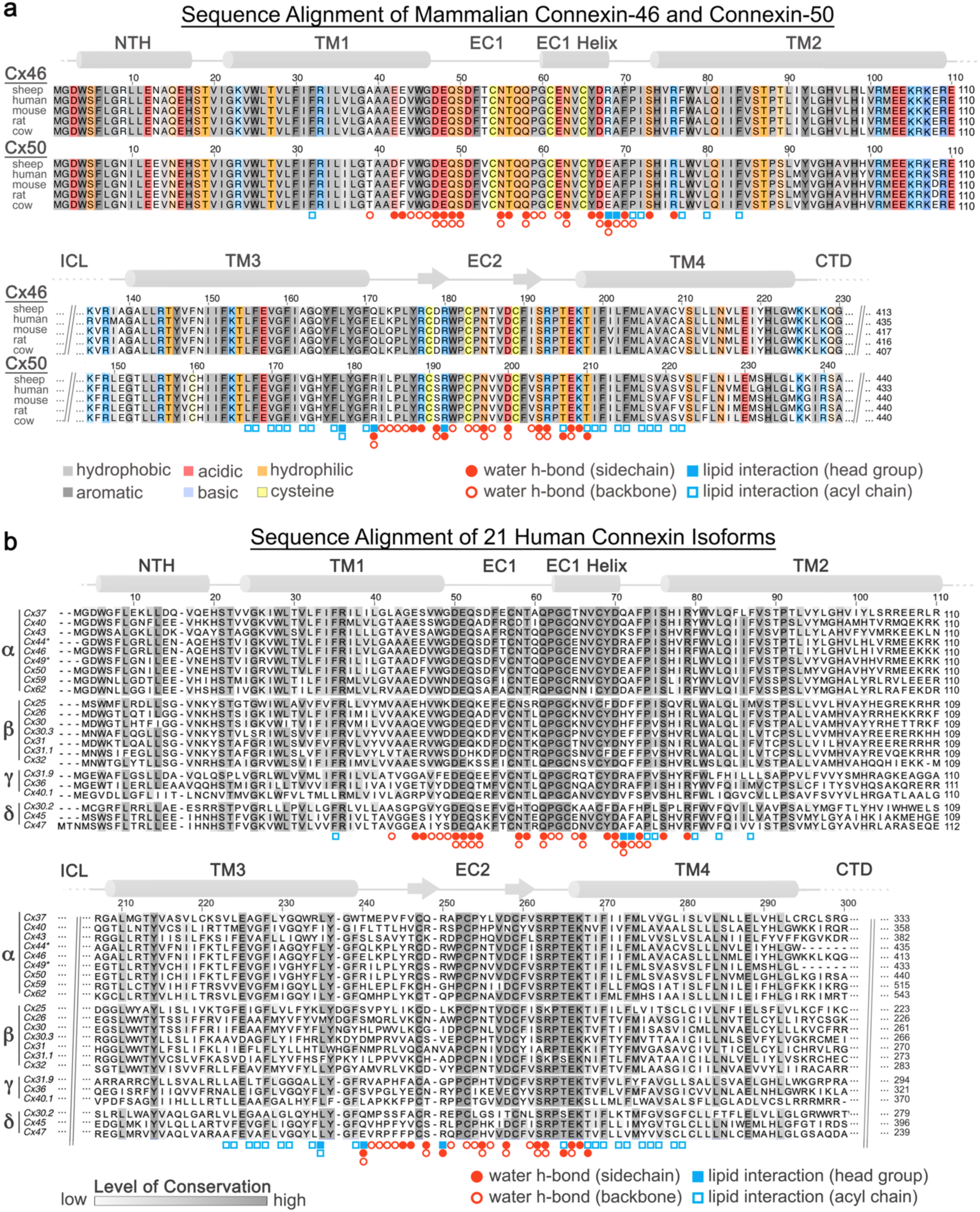
Sequence alignment with annotated lipid and water binding sites. **a)** Multiple sequence alignment of mammalian Cx46 and Cx50 isoforms with residues contributing to lipid and/or water binding sites annotated (filled circle – water h-bonding with amino acid sidechain; open circle – water h-bonding with amino acid backbone) and (filled square – interaction involving lipid headgroup; open square – interaction involving lipid acyl chain). Primary sequence coloring corresponds to amino acid type (grey – hydrophobic; dark grey – aromatic; red – acidic; blue – basic; orange – hydrophilic; yellow – cysteine). Regions of sequence homology are indicated by the level of shading. Secondary structure and domain labels are indicated for the n-terminal helix (NTH), transmembrane helices (TM1-4) and extracellular domains (EC1-2). Regions lacking defined structure and of poor sequence homology within the intracellular loop (ICL) and c-terminal domain (CTD) have been omitted for clarity. Sheep and human Cx46 and Cx50 orthologs contain ∼95% sequence identity (∼98% similarity) over the structured regions of the protein. Numbering corresponds to the amino acid sequence of sheep Cx44 and Cx49 used in the main text. **b)** Multiple sequence alignment of 20 human connexin isoforms, with sheep Cx44 (Cx46 homolog) and Cx49 (Cx50 homolog) included for comparison. Isoforms are categorized by connexin family α, β, γ and δ. The orphan Cx23 was excluded from analysis. Regions of sequence homology are indicated by the level grey of shading. Annotations for lipid and water binding sites and secondary structural elements/domains are indicated as in panel a.

**Extended Data Figure 7.**
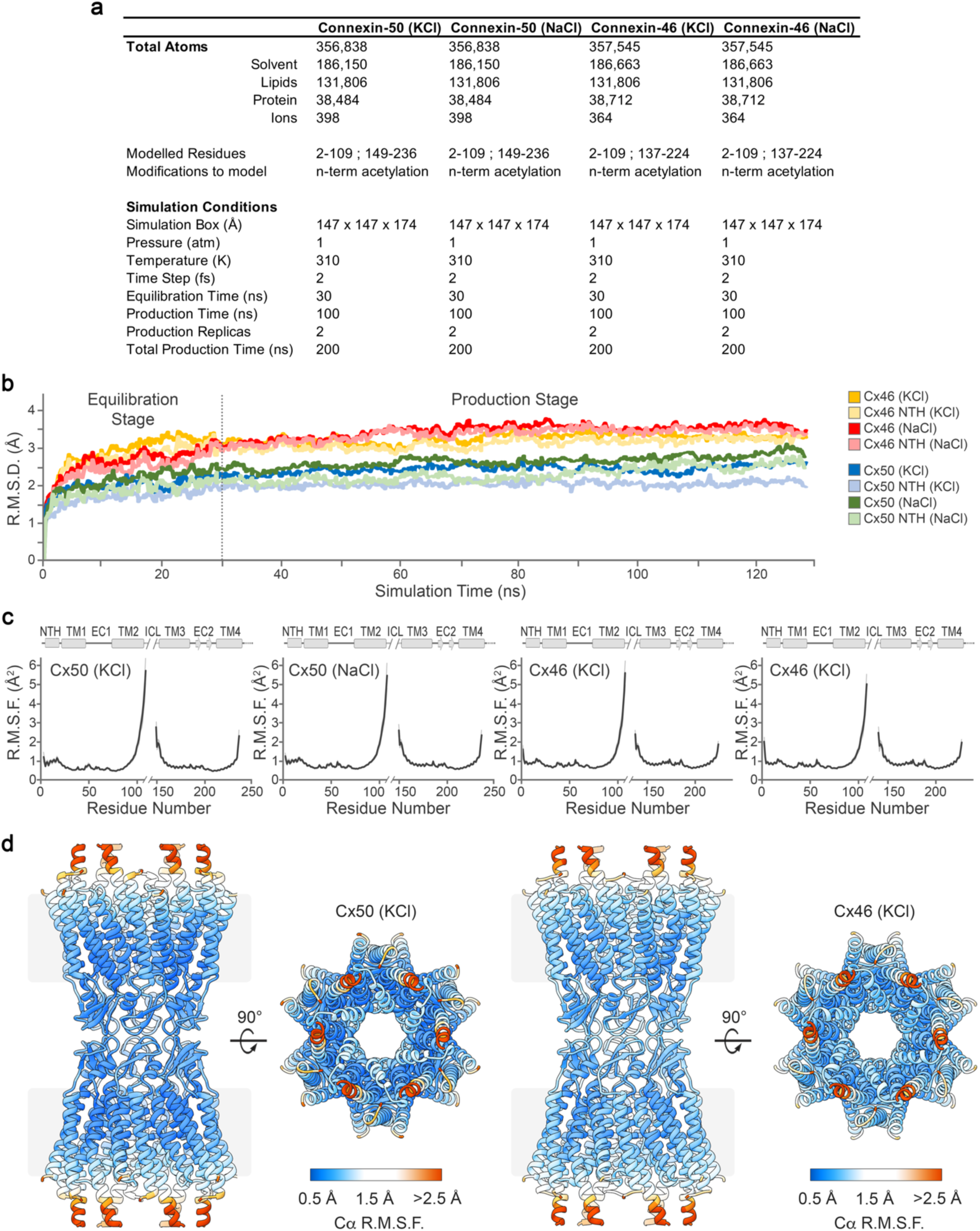
Molecular dynamics setup and validation. **a)** Summary of molecular dynamics (MD) simulation setup and conditions. Each simulation was setup similarly, using and explicit solvent model containing either KCl (Cx50KCl; Cx46 KCl) or NaCl (Cx50 NaCl; Cx46 NaCl) in the cytoplasmic space, to match either cellular or *in vitro* conditions used for CryoEM studies, respectively. All simulations were conducted with NaCl in the extracellular space and using DMPC as the lipid system. Following minimization, all systems were equilibrated for 30 ns at 37° C, and multiple replicates (N=2) of production (100 ns each) were acquired for analysis at 37° C. **b)** Cα root mean squared deviation (r.m.s.d.) analysis of equilibrium (0 – 30 ns) and production phases (30–130 ns) of the MD simulations, calculated with respect to the experimental starting structures, where Cx50 KCl (blue traces); Cx50 NaCl (green traces); Cx46 KCl (orange traces); Cx46 NaCl (red traces). Separate analysis for the n-terminal helix (NTH) domains are shown in lighter shades. **c)** Plot of average Cα root mean squared fluctuation (*r*.*m*.*s*.*f*.) during the production phase of the molecular dynamics (MD) simulations for Cx50 KCl (*left*), Cx50 NaCl *(left center)* Cx46 KCl *(right center)* and Cx46 NaCl *(right)*. Averages are determined for the 12 subunits composing the intercellular channel, analyzed for both independent productions. Error bars (light grey shading) represent 95% confidence intervals (n = 24). Secondary structure and domain labels are indicated for the n-terminal helix (NTH), transmembrane helices (TM1-4), extracellular domains (EC1-2) and intracellular loop (ICL; not modeled). **d**) Representative r.m.s.f. values mapped to the experimental starting structures of Cx50 KCl *(left)* and Cx46 KCl *(right)*. Colors correspond to r.m.s.f. amplitudes: < 0.5 Å (blue) – 1.5 Å (white) – 2.5 Å (red).

**Extended Data Figure 8.**
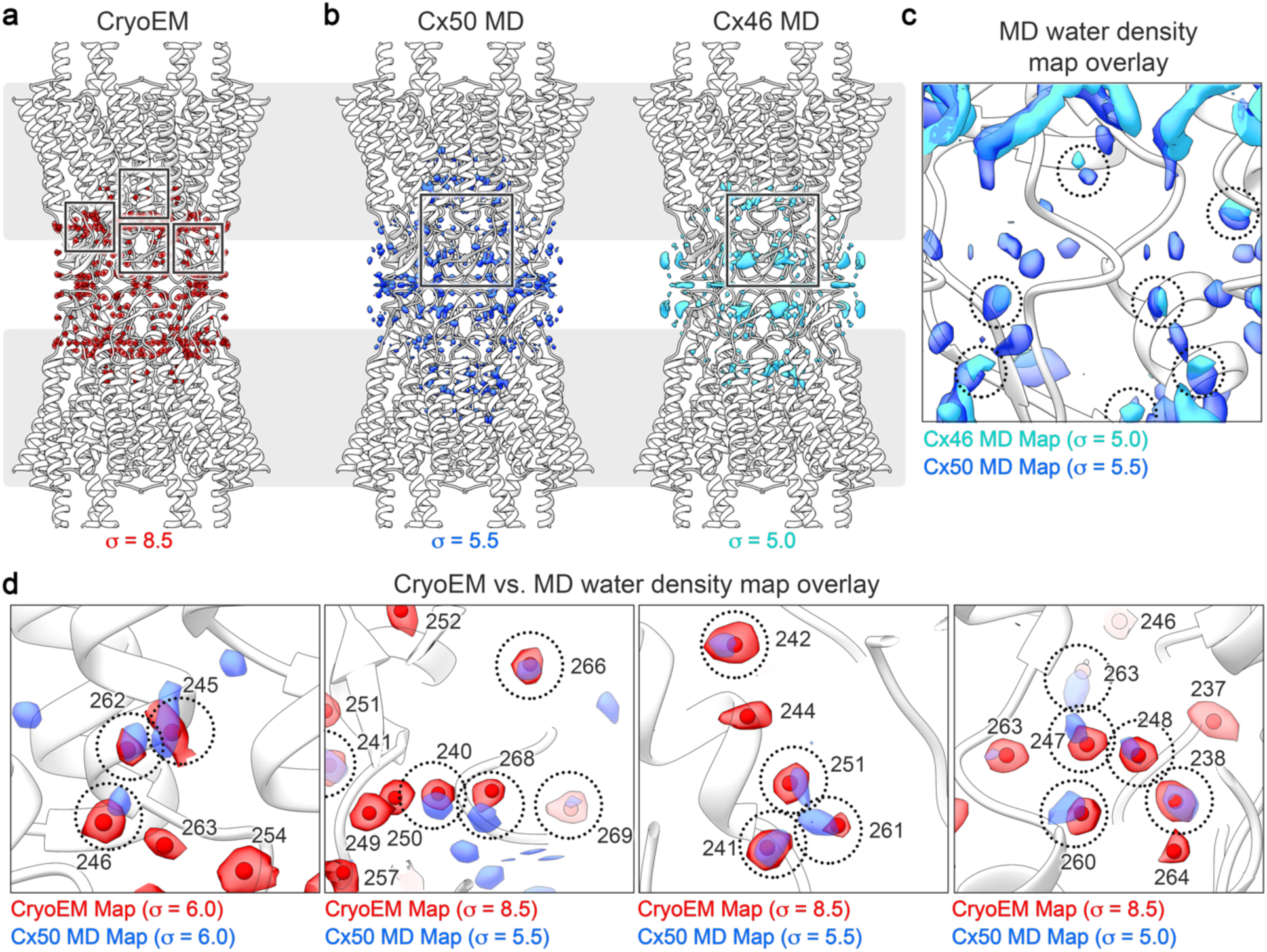
Analysis of MD-based water density maps. **a)** Ribbon structure of Cx46/50 with segmented water densities from the ensemble CryoEM map (red density, threshold = 8.5 σ). **b)** Ribbon structures of Cx50/46 with overlaid time-averaged and symmetrized water density maps calculated from MD-simulation for Cx50 (*left;* blue density, threshold = 5.5 σ) and Cx46 (*right;* cyan density, threshold = 5.0 σ). **c)** Zoom view, corresponding to boxed regions in panels b and c, showing overlaid MD-based water densities. Representative regions of overlapping density are circled. **d)** Zoom views of boxed regions in panel a, showing representative regions of CryoEM water densities (red) overlaid the Cx50 MD-based water density map (blue). Identities of modeled waters are indicated (using Cx50 numbering). Representative regions of overlapping density are circled. 76% of waters modeled into the CryoEM map show corresponding density in the MD-based water maps. Density map threshold values (σ) used for visualization in each panel are indicated.

**Extended Data Figure 9.**
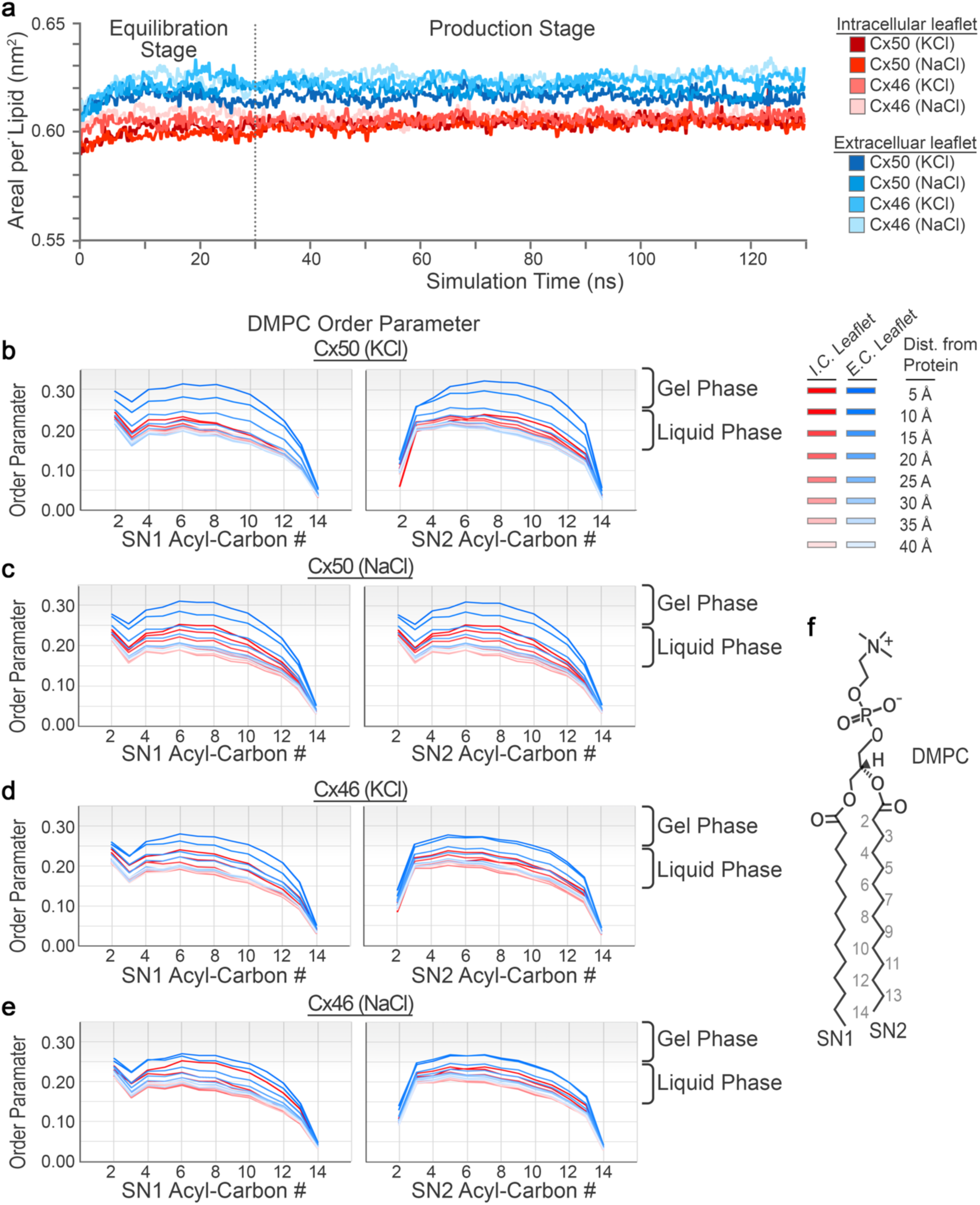
Analysis of MD-based lipid dynamics. **a)** Lipid equilibration was monitored by analyzing the averaged area per lipid (nm^2^) over the duration of MD-simulation. Traces correspond to lipids from the extracellular leaflets (blue shades) and intracellular leaflets (red shades) for each system (Cx50 KCl, Cx50 NaCl, Cx46 KCl and Cx46 NaCl) are displayed. **b-e)** Averaged lipid order parameters calculated for the SN1 *(left)* and SN2 *(right)* acyl-chain C-H bonds (S_CD_) for each system (*panel b*, Cx50 KCl; *panel c*, Cx50 NaCl; *panel d*, Cx46 KCl; and *panel e*, Cx46 NaCl). Traces correspond to lipids from the intracellular leaflets (red, *I*.*C. leaflet*) and extracellular leaflets (blue, *E*.*C. leaflet*), with dark to light shading showing the radial distance dependence from the surface of the protein (5 Å shells). **f)** Structure of dimyristoyl phosphatidylcholine (DMPC) with SN1 and SN2 acyl-chains labeled.

## Supplemental Movie Legends

**Supplemental Movie 1. CryoEM map of Cx46/50 in a dual lipid nanodisc system resolved at 1.9 Å resolution**. The CryoEM map has been segmented and colored with Cx46/50 (white), resolved lipid acyl-chains (blue) and water molecules (red). *Inset*, shows a zoom-view of the CryoEM map (transparent) with fitted atomic model of Cx50 (stick representation).

**Supplemental Movie 2. Model of Cx46/50 intercellular channel with associated lipids and water molecules**. Cx46/50 is displayed (cylinder representation) with lipid acyl-chains (blue and white spheres) and ordered water molecules displayed (red spheres).

**Supplemental Movie 3. Model of Cx46/50 monomer with associated lipids and water molecules**. Cx46/50 is displayed (cylinder representation) with 15 lipid acyl-chains (blue and white spheres) and 33 ordered water molecules displayed (red spheres).

**Supplemental Movie 4. Super-positioning of stabilized annular lipids observed by MD-simulation**. MD-trajectory, showing an ensemble super-positioning of symmetry-related DMPC lipids (displayed as all atom representation) occupying the MD-based lipid acyl-chain density map (blue). Cx46/50 is displayed in ribbon representation (white) and held static for visualization purposes.

**Supplemental Movie 5. Representative examples of stable PC lipid configurations classified from MD-simulation**. MD-trajectories, showing representative examples of DMPC lipids displayed as all atom representation (blue) occupying the MD-based lipid acyl-chain density map (grey mesh) that were classified as occupying stable configurational states (labeled, as indicated in Main Fig. 4c,d).

**Supplemental Movie 6. Representative examples of transitioning PC lipid configurations classified from MD-simulation**. MD-trajectories, showing representative examples of DMPC lipids displayed as all atom representation (blue) occupying the MD-based lipid acyl-chain density map (grey mesh) that were classified as transitioning between multiple configurational states (labeled; as indicated in Main Fig. 4c,d).

## Main Text References

1 Goodenough, D. A. & Paul, D. L. Gap junctions. Cold Spring Harbor perspectives in biology 1, a002576, doi:10.1101/cshperspect.a002576 (2009).

2 Malewicz, B., Kumar, V. V., Johnson, R. G. & Baumann, W. J. Lipids in gap junction assembly and function. Lipids 25, 419–427, doi:10.1007/bf02538083 (1990).

3 Cascio, M. Connexins and their environment: effects of lipids composition on ion channels. Biochim Biophys Acta 1711, 142–153, doi:10.1016/j.bbamem.2004.12.001 (2005).

4 Puebla, C., Retamal, M. A., Acuna, R. & Saez, J. C. Regulation of Connexin-Based Channels by Fatty Acids. Front Physiol 8, 11, doi:10.3389/fphys.2017.00011 (2017).

5 Sosinsky, G. E. & Nicholson, B. J. Structural organization of gap junction channels. Biochim Biophys Acta 1711, 99–125, doi:10.1016/j.bbamem.2005.04.001 (2005).

6 Sohl, G. & Willecke, K. Gap junctions and the connexin protein family. Cardiovasc Res 62, 228–232, doi:10.1016/j.cardiores.2003.11.013 (2004).

7 Harris, A. L. Connexin channel permeability to cytoplasmic molecules. Prog Biophys Mol Biol 94, 120–143, doi:10.1016/j.pbiomolbio.2007.03.011 (2007).

8 Bonacquisti, E. E. & Nguyen, J. Connexin 43 (C×43) in cancer: Implications for therapeutic approaches via gap junctions. Cancer Lett 442, 439–444, doi:10.1016/j.canlet.2018.10.043 (2019).

9 Delmar, M. et al. Connexins and Disease. Cold Spring Harbor perspectives in biology, doi:10.1101/cshperspect.a029348 (2017).

10 Garcia, I. E. et al. Connexinopathies: a structural and functional glimpse. BMC Cell Biol 17 Suppl 1, 17, doi:10.1186/s12860-016-0092-x (2016).

11 Aasen, T., Mesnil, M., Naus, C. C., Lampe, P. D. & Laird, D. W. Gap junctions and cancer: communicating for 50 years. Nat Rev Cancer 16, 775–788, doi:10.1038/nrc.2016.105 (2016).

12 Makowski, L., Caspar, D. L., Phillips, W. C. & Goodenough, D. A. Gap junction structures. II. Analysis of the x-ray diffraction data. J Cell Biol 74, 629–645 (1977).

13 Revel, J. P. & Karnovsky, M. J. Hexagonal array of subunits in intercellular junctions of the mouse heart and liver. J Cell Biol 33, C7–C12, doi:10.1083/jcb.33.3.c7 (1967).

14 Kistler, J., Goldie, K., Donaldson, P. & Engel, A. Reconstitution of native-type noncrystalline lens fiber gap junctions from isolated hemichannels. J Cell Biol 126, 1047–1058, doi:10.1083/jcb.126.4.1047 (1994).

15 Locke, D. & Harris, A. L. Connexin channels and phospholipids: association and modulation. BMC Biol 7, 52, doi:10.1186/1741-7007-7-52 (2009).

16 Myers, J. B. et al. Structure of native lens connexin 46/50 intercellular channels by cryo-EM. Nature 564, 372–377, doi:10.1038/s41586-018-0786-7 (2018).

17 Denisov, I. G., Grinkova, Y. V., Lazarides, A. A. & Sligar, S. G. Directed self-assembly of monodisperse phospholipid bilayer Nanodiscs with controlled size. J Am Chem Soc 126, 3477–3487, doi:10.1021/ja0393574 (2004).

18 Maeda, S. et al. Structure of the connexin 26 gap junction channel at 3.5 A resolution. Nature 458, 597–602, doi:10.1038/nature07869 (2009).

19 Bennett, B. C. et al. An electrostatic mechanism for Ca(2+)-mediated regulation of gap junction channels. Nat Commun 7, 8770, doi:10.1038/ncomms9770 (2016).

20 Gong, X. Q. & Nicholson, B. J. Size selectivity between gap junction channels composed of different connexins. Cell Commun Adhes 8, 187–192 (2001).

21 Trexler, E. B., Bukauskas, F. F., Kronengold, J., Bargiello, T. A. & Verselis, V. K. The first extracellular loop domain is a major determinant of charge selectivity in connexin46 channels. Biophys J 79, 3036–3051, doi:10.1016/S0006-3495(00)76539-8 (2000).

22 Kronengold, J., Trexler, E. B., Bukauskas, F. F., Bargiello, T. A. & Verselis, V. K. Porelining residues identified by single channel SCAM studies in Cx46 hemichannels. Cell Commun Adhes 10, 193–199 (2003).

23 Verselis, V. K., Trelles, M. P., Rubinos, C., Bargiello, T. A. & Srinivas, M. Loop gating of connexin hemichannels involves movement of pore-lining residues in the first extracellular loop domain. The Journal of biological chemistry 284, 4484–4493, doi:10.1074/jbc.M807430200 (2009).

24 Oh, S., Verselis, V. K. & Bargiello, T. A. Charges dispersed over the permeation pathway determine the charge selectivity and conductance of a C×32 chimeric hemichannel. J Physiol 586, 2445–2461, doi:10.1113/jphysiol.2008.150805 (2008).

25 Kwon, T. et al. Molecular dynamics simulations of the C×26 hemichannel: insights into voltage-dependent loop-gating. Biophys J 102, 1341–1351, doi:10.1016/j.bpj.2012.02.009 (2012).

26 Kwon, T., Harris, A. L., Rossi, A. & Bargiello, T. A. Molecular dynamics simulations of the C×26 hemichannel: evaluation of structural models with Brownian dynamics. J Gen Physiol 138, 475–493, doi:10.1085/jgp.201110679 (2011).

27 Zonta, F., Polles, G., Zanotti, G. & Mammano, F. Permeation pathway of homomeric connexin 26 and connexin 30 channels investigated by molecular dynamics. J Biomol Struct Dyn 29, 985–998, doi:10.1080/073911012010525027 (2012).

28 Bargiello, T. A., Tang, Q., Oh, S. & Kwon, T. Voltage-dependent conformational changes in connexin channels. Biochim Biophys Acta 1818, 1807–1822, doi:10.1016/j.bbamem.2011.09.019 (2012).

29 Tong, X. et al. The First Extracellular Domain Plays an Important Role in Unitary Channel Conductance of C×50 Gap Junction Channels. PLoS One 10, e0143876, doi:10.1371/journal.pone.0143876 (2015).

30 Lopez, W. et al. Mechanism of gating by calcium in connexin hemichannels. Proc Natl Acad Sci U S A 113, E7986–E7995, doi:10.1073/pnas.1609378113 (2016).

31 Garcia, I. E. et al. The syndromic deafness mutation G12R impairs fast and slow gating in C×26 hemichannels. J Gen Physiol 150, 697–711, doi:10.1085/jgp.201711782 (2018).

32 Rubinos, C., Sanchez, H. A., Verselis, V. K. & Srinivas, M. Mechanism of inhibition of connexin channels by the quinine derivative N-benzylquininium. J Gen Physiol 139, 69–82, doi:10.1085/jgp.201110678 (2012).

33 Banks, E. A. et al. Connexin mutation that causes dominant congenital cataracts inhibits gap junctions, but not hemichannels, in a dominant negative manner. J Cell Sci 122, 378–388, doi:10.1242/jcs.034124 (2009).

34 Berthoud, V. M. et al. Connexin50D47A decreases levels of fiber cell connexins and impairs lens fiber cell differentiation. Invest Ophthalmol Vis Sci 54, 7614–7622, doi:10.1167/iovs.13-13188 (2013).

35 Reis, L. M. et al. Whole exome sequencing in dominant cataract identifies a new causative factor, CRYBA2, and a variety of novel alleles in known genes. Hum Genet 132, 761–770, doi:10.1007/s00439-013-1289-0 (2013).

36 White, T. W., Bruzzone, R., Wolfram, S., Paul, D. L. & Goodenough, D. A. Selective interactions among the multiple connexin proteins expressed in the vertebrate lens: the second extracellular domain is a determinant of compatibility between connexins. J Cell Biol 125, 879–892 (1994).

37 White, T. W., Paul, D. L., Goodenough, D. A. & Bruzzone, R. Functional analysis of selective interactions among rodent connexins. Mol Biol Cell 6, 459–470, doi:10.1091/mbc.6.4.459 (1995).

38 Nakagawa, S. et al. Asparagine 175 of connexin32 is a critical residue for docking and forming functional heterotypic gap junction channels with connexin26. The Journal of biological chemistry 286, 19672–19681, doi:10.1074/jbc.M110.204958 (2011).

39 Cottrell, G. T. & Burt, J. M. Functional consequences of heterogeneous gap junction channel formation and its influence in health and disease. Biochim Biophys Acta 1711, 126–141, doi:10.1016/j.bbamem.2004.11.013 (2005).

40 Bai, D. & Wang, A. H. Extracellular domains play different roles in gap junction formation and docking compatibility. Biochem J 458, 1–10, doi:10.1042/BJ20131162 (2014).

41 Schadzek, P. et al. The cataract related mutation N188T in human connexin46 (hC×46) revealed a critical role for residue N188 in the docking process of gap junction channels. Biochim Biophys Acta 1858, 57–66, doi:10.1016/j.bbamem.2015.10.001 (2016).

42 Silander, K. et al. Spectrum of mutations in Finnish patients with Charcot-Marie-Tooth disease and related neuropathies. Hum Mutat 12, 59–68, doi:10.1002/(SICI)1098-1004(1998)12:1<59::AID-HUMU9>3.0.CO;2-A (1998).

43 Primignani, P. et al. A novel dominant missense mutation--D179N--in the GJB2 gene (Connexin 26) associated with non-syndromic hearing loss. Clin Genet 63, 516–521, doi:10.1034/j.1399-0004.2003.00079.x (2003).

44 Deeley, J. M. et al. Human lens lipids differ markedly from those of commonly used experimental animals. Biochim Biophys Acta 1781, 288–298, doi:10.1016/j.bbalip.2008.04.002 (2008).

45 Lampe, P. D. et al. In vitro assembly of gap junctions. J Struct Biol 107, 281–290 (1991).

46 Mabrey, S. & Sturtevant, J. M. Investigation of phase transitions of lipids and lipid mixtures by sensitivity differential scanning calorimetry. Proc Natl Acad Sci U S A 73, 3862–3866, doi:10.1073/pnas.73.11.3862 (1976).

47 Shaw, A. W., McLean, M. A. & Sligar, S. G. Phospholipid phase transitions in homogeneous nanometer scale bilayer discs. FEBS Lett 556, 260–264, doi:10.1016/s0014-5793(03)01400-5 (2004).

48 Vermeer, L. S., de Groot, B. L., Reat, V., Milon, A. & Czaplicki, J. Acyl chain order parameter profiles in phospholipid bilayers: computation from molecular dynamics simulations and comparison with 2H NMR experiments. Eur Biophys J 36, 919–931, doi:10.1007/s00249-007-0192-9 (2007).

49 Khakbaz, P. & Klauda, J. B. Investigation of phase transitions of saturated phosphocholine lipid bilayers via molecular dynamics simulations. Biochim Biophys Acta Biomembr 1860, 1489–1501, doi:10.1016/j.bbamem.2018.04.014 (2018).

50 Caspar, D. L., Goodenough, D. A., Makowski, L. & Phillips, W. C. Gap junction structures. Correlated electron microscopy and x-ray diffraction. J Cell Biol 74, 605–628 (1977).

51 Schubert, A. L., Schubert, W., Spray, D. C. & Lisanti, M. P. Connexin family members target to lipid raft domains and interact with caveolin-1. Biochemistry 41, 5754–5764, doi:10.1021/bi0121656 (2002).

52 Locke, D., Liu, J. & Harris, A. L. Lipid rafts prepared by different methods contain different connexin channels, but gap junctions are not lipid rafts. Biochemistry 44, 13027–13042, doi:10.1021/bi050495a (2005).

53 Hunte, C. Specific protein-lipid interactions in membrane proteins. Biochem Soc Trans 33, 938–942 (2005).

54 Spray, D. C., Rozental, R. & Srinivas, M. Prospects for rational development of pharmacological gap junction channel blockers. Current drug targets 3, 455–464 (2002).

## Additional References

55 Ritchie, T. K. et al. Chapter 11 - Reconstitution of membrane proteins in phospholipid bilayer nanodiscs. Methods Enzymol 464, 211–231, doi:10.1016/S0076-6879(09)64011-8 (2009).

56 Kistler, J., Christie, D. & Bullivant, S. Homologies between gap junction proteins in lens, heart and liver. Nature 331, 721–723, doi:10.1038/331721a0 (1988).

57 Kistler, J., Schaller, J. & Sigrist, H. MP38 contains the membrane-embedded domain of the lens fiber gap junction protein MP70. The Journal of biological chemistry 265, 13357–13361 (1990).

58 White, T. W., Bruzzone, R., Goodenough, D. A. & Paul, D. L. Mouse Cx50, a functional member of the connexin family of gap junction proteins, is the lens fiber protein MP70. Mol Biol Cell 3, 711–720 (1992).

59 Wang, Z. & Schey, K. L. Phosphorylation and truncation sites of bovine lens connexin 46 and connexin 50. Exp Eye Res 89, 898–904, doi:10.1016/j.exer.2009.07.015 (2009).

60 Reichow, S. L. et al. Allosteric mechanism of water-channel gating by Ca2+-calmodulin. Nat Struct Mol Biol 20, 1085–1092, doi:10.1038/nsmb.2630 (2013).

61 Gold, M. G. et al. AKAP2 anchors PKA with aquaporin-0 to support ocular lens transparency. EMBO Mol Med 4, 15–26, doi:10.1002/emmm.201100184 (2012).

62 Reichow, S. L. & Gonen, T. Noncanonical binding of calmodulin to aquaporin-0: implications for channel regulation. Structure 16, 1389–1398, doi:10.1016/j.str.2008.06.011 (2008).

63 Efremov, R. G., Gatsogiannis, C. & Raunser, S. Lipid Nanodiscs as a Tool for High-Resolution Structure Determination of Membrane Proteins by Single-Particle Cryo-EM. Methods Enzymol 594, 1–30, doi:10.1016/bs.mie.2017.05.007 (2017).

64 Tang, G. et al. EMAN2: an extensible image processing suite for electron microscopy. J Struct Biol 157, 38–46, doi:10.1016/j.jsb.2006.05.009 (2007).

65 Ludtke, S. J. Single-Particle Refinement and Variability Analysis in EMAN2.1. Methods Enzymol 579, 159–189, doi:10.1016/bs.mie.2016.05.001 (2016).

66 Zivanov, J. et al. New tools for automated high-resolution cryo-EM structure determination in RELION-3. eLife 7, doi:10.7554/eLife.42166 (2018).

67 Zhang, K. Gctf: Real-time CTF determination and correction. J Struct Biol 193, 1–12, doi:10.1016/j.jsb.2015.11.003 (2016).

68 Zivanov, J., Nakane, T. & Scheres, S. H. W. Estimation of high-order aberrations and anisotropic magnification from cryo-EM data sets in RELION-3.1. IUCrJ 7, 253–267, doi:10.1107/S2052252520000081 (2020).

69 Scheres, S. H. & Chen, S. Prevention of overfitting in cryo-EM structure determination. Nat Methods 9, 853–854, doi:10.1038/nmeth.2115 (2012).

70 Pettersen, E. F. et al. UCSF Chimera--a visualization system for exploratory research and analysis. J Comput Chem 25, 1605–1612, doi:10.1002/jcc.20084 (2004).

71 Emsley, P., Lohkamp, B., Scott, W. G. & Cowtan, K. Features and development of Coot. Acta Crystallogr. D66, 486–501 (2010).

72 Afonine, P. V. et al. Real-space refinement in PHENIX for cryo-EM and crystallography. Acta Crystallogr D Struct Biol 74, 531–544, doi:10.1107/S2059798318006551 (2018).

73 Williams, C. J. et al. MolProbity: More and better reference data for improved all-atom structure validation. Protein Sci 27, 293–315, doi:10.1002/pro.3330 (2018).

74 Barad, B. A. et al. EMRinger: side chain-directed model and map validation for 3D cryo-electron microscopy. Nat Methods 12, 943–946, doi:10.1038/nmeth.3541 (2015).

75 Moriarty, N. W., Grosse-Kunstleve, R. W. & Adams, P. D. electronic Ligand Builder and Optimization Workbench (eLBOW): a tool for ligand coordinate and restraint generation. Acta Crystallogr D Biol Crystallogr 65, 1074–1080, doi:10.1107/S0907444909029436 (2009).

76 Konig, N. & Zampighi, G. A. Purification of bovine lens cell-to-cell channels composed of connexin44 and connexin50. J Cell Sci 108 (Pt 9), 3091–3098 (1995).

77 Jiang, J. X. & Goodenough, D. A. Heteromeric connexons in lens gap junction channels. Proc Natl Acad Sci U S A 93, 1287–1291 (1996).

78 Humphrey, W., Dalke, A. & Schulten, K. VMD: visual molecular dynamics. J Mol Graph 14, 33–38, 27-38 (1996).

79 Shearer, D., Ens, W., Standing, K. & Valdimarsson, G. Posttranslational modifications in lens fiber connexins identified by off-line-HPLC MALDI-quadrupole time-of-flight mass spectrometry. Invest Ophthalmol Vis Sci 49, 1553–1562, doi:10.1167/iovs.07-1193 (2008).

80 Varland, S., Osberg, C. & Arnesen, T. N-terminal modifications of cellular proteins: The enzymes involved, their substrate specificities and biological effects. Proteomics 15, 2385–2401, doi:10.1002/pmic.201400619 (2015).

81 Grubmuller, H., Heymann, B. & Tavan, P. Ligand binding: molecular mechanics calculation of the streptavidin-biotin rupture force. Science 271, 997–999 (1996).

82 Wu, E. L. et al. CHARMM-GUI Membrane Builder toward realistic biological membrane simulations. J Comput Chem 35, 1997–2004, doi:10.1002/jcc.23702 (2014).

83 Phillips, J. C. et al. Scalable molecular dynamics with NAMD. J Comput Chem 26, 1781–1802, doi:10.1002/jcc.20289 (2005).

84 Huang, J. & MacKerell, A. D., Jr. CHARMM36 all-atom additive protein force field: validation based on comparison to NMR data. J Comput Chem 34, 2135–2145, doi:10.1002/jcc.23354 (2013).

85 Buchoux, S. FATSLiM: a fast and robust software to analyze MD simulations of membranes. Bioinformatics 33, 133–134, doi:10.1093/bioinformatics/btw563 (2017).

86 Piggot, T. J., Allison, J. R., Sessions, R. B. & Essex, J. W. On the Calculation of Acyl Chain Order Parameters from Lipid Simulations. J Chem Theory Comput 13, 5683–5696, doi:10.1021/acs.jctc.7b00643 (2017).

87 Donaldson, P. & Kistler, J. Reconstitution of channels from preparations enriched in lens gap junction protein MP70. J Membr Biol 129, 155–165 (1992).

88 Smart, O. S., Neduvelil, J. G., Wang, X., Wallace, B. A. & Sansom, M. S. HOLE: a program for the analysis of the pore dimensions of ion channel structural models. J Mol Graph 14, 354–360, 376 (1996).

